# *MIR319C* activation by CUP-SHAPED COTYLEDON2 delays cell proliferation-to-differentiation transition in Arabidopsis leaf primordia

**DOI:** 10.1101/2024.09.07.611804

**Authors:** Naveen Shankar, Abhishek Gupta, Vishwadeep Mane, Somsree Roy, Ahana Jayanarayan Perumadaraya, V Avani, Olivier Hamant, Utpal Nath

## Abstract

The microRNA miR319 regulates leaf size in diverse plant species by reducing the level of the target transcripts that encode JAW-TCPs, the transcription factors (TF) that restrict leaf size by committing the proliferating pavement cells to differentiation. *MIR319C*, one of the three miR319-producing genes in Arabidopsis, is expressed throughout the incipient leaf primordia, and its expression domain gets restricted to the base at later stages, partly due to its transcriptional repression by JAW-TCPs. However, the factors that activate and maintain *MIR319C* expression in leaf primordia are yet unknown. Here, we identify the CUP-SHAPED COTYLEDON2 (CUC2) TF as a direct activator of *MIR319C* transcription. Using a yeast one-hybrid (Y1H) screen, we identified several NAC domain TFs as potential regulators of *MIR319C*. Subsequent ex vivo binding and transactivation assays suggested that CUC2 binds to a distal promoter region of the *MIR319C* locus and activates its transcription. Mutants with compromised *CUC2* and *MIR319C* activities resulted in smaller leaves with fewer cells. Detailed morphometric analysis of higher order *CUC2* and *MIR319* loss-of-function mutants highlighted the crucial role of the *CUC2-MIR319* module in maintaining the duration of cell proliferation in leaf primordia. Additionally, the phenotype of mutants with altered *CUC2* and *MIR319/JAW-TCP* activities demonstrated that *CUC2* enhances leaf size through the *MIR319C-JAW-TCP* pathway. Overall, our findings uncovered a novel role for CUC2 in sustaining cell division by activating *MIR319C* transcription in the leaf primordia.

## Introduction

Leaf development can be empirically divided into three major stages namely initiation, primary morphogenesis, and secondary morphogenesis (Bar and Ori 2014). Initiation of a leaf primordium commences with the specification of a group of founder cells at the periphery of the shoot apical meristem (SAM), a dome-shaped structure at the shoot apex that harbours a constant pool of stem cells throughout plant life (Barton 2010). Concurrently, a low cell proliferation region, termed as the ‘boundary zone’, gets established at the interface that separates the primordium from the SAM (Hepworth and Lenhard 2014; Wang et al. 2016). The basic form of the leaf consisting of the lamina and the petiole is established in eudicots during primary morphogenesis, wherein leaf growth is primarily sustained by cell proliferation. The end of primary morphogenesis is marked by the switch from cell proliferation to differentiation and the formation of specific cell types such as trichomes and stomata (Bar and Ori 2014). During secondary morphogenesis, differentiation of all the cell types is completed, and the leaf grows primarily by cell expansion (Bar and Ori 2014; Du et al. 2018; Maugarny-Calès and Laufs 2018).

The shape and size of the mature leaf are regulated at each of the three developmental stages by modulating the number of founder cells during initiation, and by regulating the rate, location and duration of cell proliferation and expansion (Gonzalez et al. 2012; Hepworth and Lenhard 2014; Kalve et al. 2014). The timing of cell proliferation arrest, which directly influences the number of cells in mature leaves, plays a critical role in determining the final leaf size across many plant species (Gázquez and Beemster 2017), suggesting that the transition from cell division to expansion is a key cellular mechanism shaped by organ size evolution. In Arabidopsis leaves, arrest of cell proliferation occurs basipetally; cells towards the tip stop dividing and commit to differentiation early (Donnelly et al. 1999; Andriankaja et al. 2012; Fox et al. 2018), while the cells towards the base continue to proliferate. Despite the critical role of the timing in the transition from cell proliferation to differentiation, its molecular basis remains less understood. Among the factors essential for this process, two microRNA-transcription factor modules control the transition from cell proliferation to cell expansion: miR396 and *GROWTH REGULATING FACTORs* (*GRFs*), and miR319 and five *CINCINNATA-*like *TEOSINTE BRANCHED 1*, *CYCLOIDEA*, *PROLIFERATING CELL FACTOR1/2* (*CIN*-*TCPs*) (Rodriguez et al. 2014; Sarvepalli and Nath 2018). miR396 downregulates seven *GRF* transcripts in the distal region of the leaf, thereby restricting GRF activities towards the base (Rodriguez et al. 2010). GRFs interact with GRF-INTERACTING FACTOR 1/ANGUSTIFOLIA 3 (GIF1/AN3) transcriptional coactivators and upregulate *CYCLIN B* (*CYCB*) expression to delay the proliferation-to-differentiation transition (Schneider et al. 2023). Eight *CIN*-*TCP* genes promote leaf maturation by triggering the transition of cell proliferation-to-differentiation in diverse eudicots including Arabidopsis (Sarvepalli and Nath 2018). The spatiotemporal pattern of *CIN-TCP* transcripts is regulated by a combined action of their promoter activity and miR319 activity (Palatnik et al. 2003; Efroni et al. 2008), which prevents the accumulation of transcripts of five *CIN-TCPs* (*TCP2, 3, 4, 10, 24*), also known as *JAGGED AND WAVY TCPs* (*JAW-TCPs*), in the proximal region of the young leaf primordia (Palatnik et al. 2003; Nag et al. 2009). Expression of miR319-resistant form of *JAW-TCPs* under endogenous regulation leads to smaller leaves with fewer cells due to precocious proliferation arrest, while ectopic expression of miR319 results in larger leaves with wavy margin due to a delay in the transition from proliferation to differentiation (Palatnik et al. 2003; Efroni et al. 2008; Challa et al. 2019). CIN-TCPs commit the leaf cells towards differentiation by modulating multiple pathways, such as auxin and cytokinin response, meristematic activity, and cell cycle progression (Sarvepalli and Nath 2018).

The *CUP SHAPED COTYLEDON* (*CUC*) group of genes, which encode the NAM/ ATAF/ CUC (NAC) group of TFs, regulate boundary development in diverse angiosperms (Wang et al. 2016). The Arabidopsis *CUC* members *CUC1*, *CUC2*, and *CUC3* are consistently expressed in the boundary zones, and their absence leads to the fusion of adjacent organs such as cotyledons, sepals, stamens, and ovules (Aida et al. 1997; Ishida et al. 2000; Vroemen et al. 2003). The *CUC* genes are also required for the maintenance of shoot meristem via the activation of the *SHOOT MERISTEMLESS* (*STM*) gene and its homologues (Aida et al. 1999; Hibara et al. 2006; Scofield et al. 2018). *CUC1* transcript is not detected in leaves, while *CUC2* and *CUC3* are broadly expressed along the SAM-leaf boundary and leaf margins (Nikovics et al. 2006; Hasson et al. 2011). However, *CUC3* shows weaker expression, and its activity is dependent on *CUC2* (Hasson et al. 2011). In leaf primordia, CUC2 and CUC3 proteins accumulate at distinct foci along the margins where they are necessary for the initiation and growth of serrations through a feedback mechanism involving auxin and the auxin efflux carrier protein PINFORMED1 (PIN1) (Nikovics et al. 2006; Bilsborough et al. 2011; Hasson et al. 2011; Maugarny-Calès et al. 2019). Interestingly, some of the gain-of-function phenotypes of *JAW-TCPs* such as impaired SAM function, cotyledon fusion, and smooth leaf margin are dependent on the downregulation of *CUC* activity (Koyama et al. 2007; Koyama et al. 2010; Koyama et al. 2017; Challa et al. 2019). While TCP3 inhibits *CUC1* and *CUC2* by activating the transcription of the miR164-encoding gene *MIR164A* to promote shoot morphogenesis (Koyama et al. 2007; Koyama et al. 2010), TCP4 interacts with CUC and inhibits their homo-dimerization and transactivation abilities to modulate serration formation and outgrowth (Rubio-Somoza et al. 2014).

Expression analysis of the miR319-encoding genes *MIR319A*, *MIR319B*, and *MIR319C* in Arabidopsis suggested their distinct developmental functions due to largely non-overlapping expression pattern (Nag et al. 2009). Whereas *MIR319A* and *MIR319B* promoter activity is not detected in leaves, *MIR319C* is strongly expressed in the leaf primordia (Nag et al. 2009; Shankar et al. 2023). The single loss-of-function mutants for miR319a and miR319b show a moderate decrease in leaf serration, and the *miR319a;miR319b* double mutant leaves lack leaf serration, suggesting a role for *MIR319A* and *MIR319B* genes in leaf development (Koyama et al. 2017). However, the *miR319a;miR319b* leaves are as big as wild-type (Koyama et al. 2017), pointing towards a key role of miR319c in downregulating *JAW-TCP* activity during the early leaf growth. We had earlier demonstrated that JAW-TCPs transcriptionally repress the miR319-encoding gene *MIR319C* in the distal region to prevent miR319 accumulation, thereby establishing spatially separated proximal miR319 domain and distal JAW-TCP domain in incipient leaf primordia (Shankar et al. 2023). However, it is still unknown how the *MIR319* genes are transcriptionally activated in early leaf primordia. Here, we demonstrate that the CUC2 protein transcriptionally activates the *MIR319C* gene and, together with the miR319 product of *MIR319A* and *MIR319B*, promotes cell proliferation and leaf size by downregulating *JAW-TCP* level.

## Results

### Transcriptional activators of *MIR319C* identified in a yeast one-hybrid screen

The *MIR319C* promoter activity is primarily detected at the shoot apex, more specifically at the meristem-primordia boundary and in young leaf primordia and is excluded from the central domain of apical meristem (Shankar et al. 2023) (**Fig. S1A, S1B**), suggesting that proteins encoded by the transcriptome subset within this domain include factors that activate *MIR319C* transcription. To identify these regulators, we screened a subset of yeast one-hybrid library containing 378 Arabidopsis transcription factors, enriched in the shoot apex, for their ability to bind to the upstream regulatory region (URR) of *MIR319C* (See methods, **Fig. S1C, S1D, Suppl Data Set. 1**). Fifty-seven TFs were identified that promoted the growth of yeast cells when a 2736 bp URR was used as bait to drive the expression of the *HISTIDINE3* (*HIS3*) selection gene (**Suppl Data Set. 2**). Gene ontology analysis revealed that TFs belonging to NAC, TCP, ETHYLENE RESPONSE FACTOR (ERF), BASIC LEUCINE ZIPPER (bZIP), and BASIC HELIX LOOP HELIX (bHLH) families were over-represented among the positive clones identified in the screen (**Fig. 1A**).

**Figure 1:**
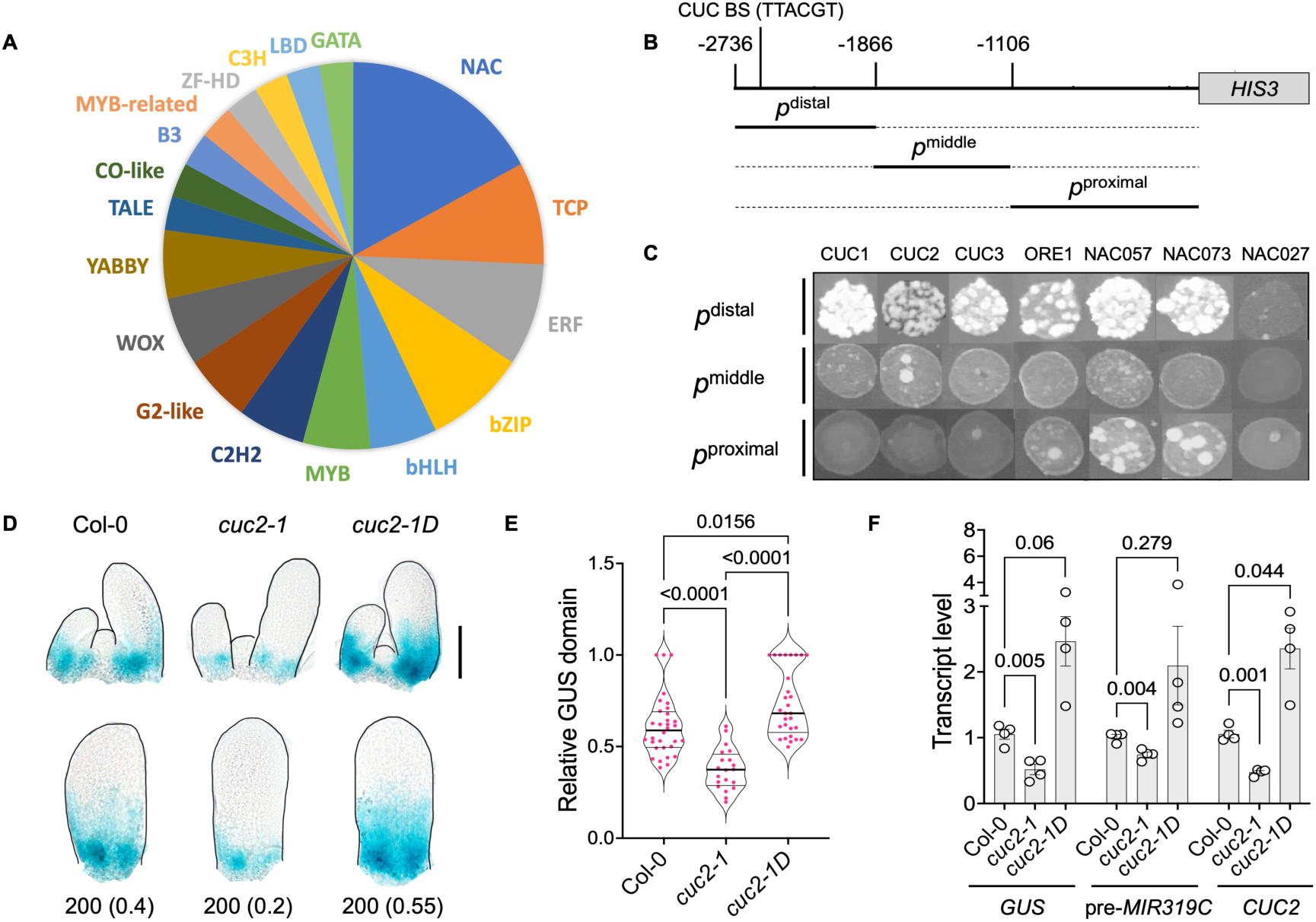
**Identification of CUC proteins as activators of *MIR319C* transcription**. **(A)** Abundance of transcription factor (TF) family members (in %) recovered from the yeast one hybrid screen (Y1H) with 378 TFs as prey proteins and 2736 bp *MIR319C* URR fragment as bait. **(B)** A schematic representation of truncated *MIR319C* promoter fragments used for Y1H assay with selected NAC TFs as prey proteins. Grey box corresponds to *HIS3 CDS*. The three bold horizontal lines below the URR scheme represent the promoter regions with truncation (dotted lines), i.e., *p*^distal^ (-2736-1866 bp), *p*^middle^ (-1866-1106 bp) and *p*^proximal^ (-1106 bp), respectively. The consensus CUC-binding sequence (BS) TTACGT is located between-2616 bp to-2621 bp positions. **(C)** Images of yeast growth harboring three different truncated *MIR319C* promoter regions upstream to the *HIS3* reporter gene and corresponding NAC TF preys as indicated, grown in the presence of inhibitory concentration of 3-AT. NAC027 served as a negative control. **(D)** Bright field images of shoot apices (top panel) or 6^th^ leaf primordia (bottom panel) of 10-day old seedlings, processed for GUS-staining, of indicated genotypes expressing *pMIR319C::GUS* transgene. Black lines highlight the boundaries. Numbers below the images in lower panel indicate leaf lengths; GUS domain length relative to the leaf length (Relative GUS domain) is indicated within parentheses. Scale bar, 100 μm. **(E)** Violin plots displaying the relative GUS domain (Y-axis) in the genotypes indicated on the X-axis. Differences among samples are indicated by *p*-values on top of the data bars; one-way ANOVA, Dunnett’s post hoc test was used. N, 20-32. **(F)** Level of *GUS,* pre-*MIR319C* and *CUC2* transcripts quantified by RT-qPCR in genotypes indicated on the X-axis. Error bars represent SEM of four biological replicates. Multiple comparisons were performed using One-way ANOVA followed by Dunnett’s post hoc test. *p*-values are shown above the data bars.

Five out of the 6 NAC prey proteins recovered from the Y1H screen (**Fig. S1E**) show comparable DNA-binding preferences to the target DNA TT(G/A)CGT (Lindemose et al. 2014), a sequence element present at the 2616-2621 nucleotide position upstream to the start site of pre-*MIR319C* sequence (**Fig. 1B**). The binding preference of these 5 NACs is also shared by other members of the NAM, OsNAC7, NAC2, ANAC011, and AtNAC3 subgroups of the AtNAC TF family (Ooka et al. 2003; Lindemose et al. 2014). To identify additional NAC members capable of binding to the *MIR319C* URR in a targeted, small-scale Y1H screen, we tested the binding ability of 20 representative members from these 5 AtNAC subgroups (**Fig. S2A, S2B**) to three non-overlapping *MIR319C* URR fragments named as *p^distal^* (-2736 to -1866 bp), *p^middle^* (-1866 to -1106 bp), and *p^proximal^* (-1106 to -1 bp) (**Fig. 1B**). Five additional NAC members including CUC1, CUC2 & CUC3 supported yeast growth when *p^distal^*, which includes the putative NAC-binding DNA element, was used as bait (**Fig. 1C)**.

To test which URR region is important for *MIR319C* promoter activity, we generated transgenic lines expressing the *GUS* reporter construct under truncated (-533 bp or -1866 bp long) or full-length (-2736 bp long) promoter fragments, and studied *pMIR319C::GUS* expression pattern in young leaves. The *MIR319C* activity was significantly weaker in the line carrying -1866 bp promoter region, and barely detectable in the line carrying -533 bp promoter region, compared to the full length promoter (**Fig. S3B**). This suggests that the distal (-2736 to -1866 bp) and the middle (-1866 to -1106 bp) promoter regions contain cis elements required for *MIR319C* expression.

### CUC2 activates *MIR319C* in leaf primordia

*CUC2*, the primary *CUC* member in leaves (Nikovics et al. 2006; Hasson et al. 2011), is active throughout the shoot apex, nascent leaf primordia, and at the base of young leaves, overlapping the *MIR319C* expression domain (**Fig. S1A, B**). As the CUC2 protein binds to *MIR319C* URR in yeast (**Fig. 1**), we explored whether CUC2 regulates *MIR319C* promoter activity in developing leaves. Comparative GUS reporter activity analysis in leaves expressing the *pMIR319C::GUS^#1^* construct (Nag et al. 2009; Shankar et al. 2023) revealed that the *MIR319C* promoter activity is altered in the *CUC2* mutants compared to wild type (**Fig. 1D, 1E**). The domain of *pMIR319C::GUS* expression relative to the leaf length was proximally restricted (median ∼0.35) in the *cuc2-1* loss-of-function leaves (Hasson et al. 2011), and distally extended (median ∼0.75) in the *cuc2-1D* gain-of-function leaves (Larue et al., 2009), compared to wild type (median ∼0.6). Corresponding to the change in the GUS domain in the mutant leaves, there was a ∼2-fold reduction in the level of *GUS* transcript in the *pMIR319C::GUS^#1^;cuc2-1* seedlings (p value, 0.005), and ∼2.5-fold increase in the *pMIR319C::GUS^#1^;cuc2-1D* seedlings (p value, 0.06), as analyzed by RT-qPCR (**Fig. 1F**), suggesting that CUC2 activates *MIR319C* promoter. This was further supported by the observation that the pre-*MIR319C* transcript level was ∼1.5-fold lower in *pMIR319C::GUS^#1^;cuc2-1* (p value, 0.004) and >2-fold higher in *pMIR319C::GUS^#1^;cuc2-1D* seedlings (p value, 0.279; **Fig. 1F**). Together, these results suggest that CUC2 is required and sufficient to activate *MIR319C* expression in leaf primordia.

### CUC2 binds to *MIR319C* locus and activates its transcription

To determine the functional significance of the predicted CUC-binding cis element within the *MIR319C* URR, we generated a GUS-based reporter line of *MIR319C* promoter carrying mutations in the CUC-binding site (**Fig. 2A**) and determined the GUS activity in the resultant *mCUCBS-pMIR319C::GUS* leaves. The relative GUS domain was much reduced in the leaf primordia expressing *mCUCBS-pMIR319C::GUS* transgene compared to the wild-type *pMIR319C::GUS^#2^* transgene (Shankar et al. 2023) (**Fig. 2B-D**), suggesting that the CUC-binding site is required for the *MIR319C* promoter activity.

**Figure 2:**
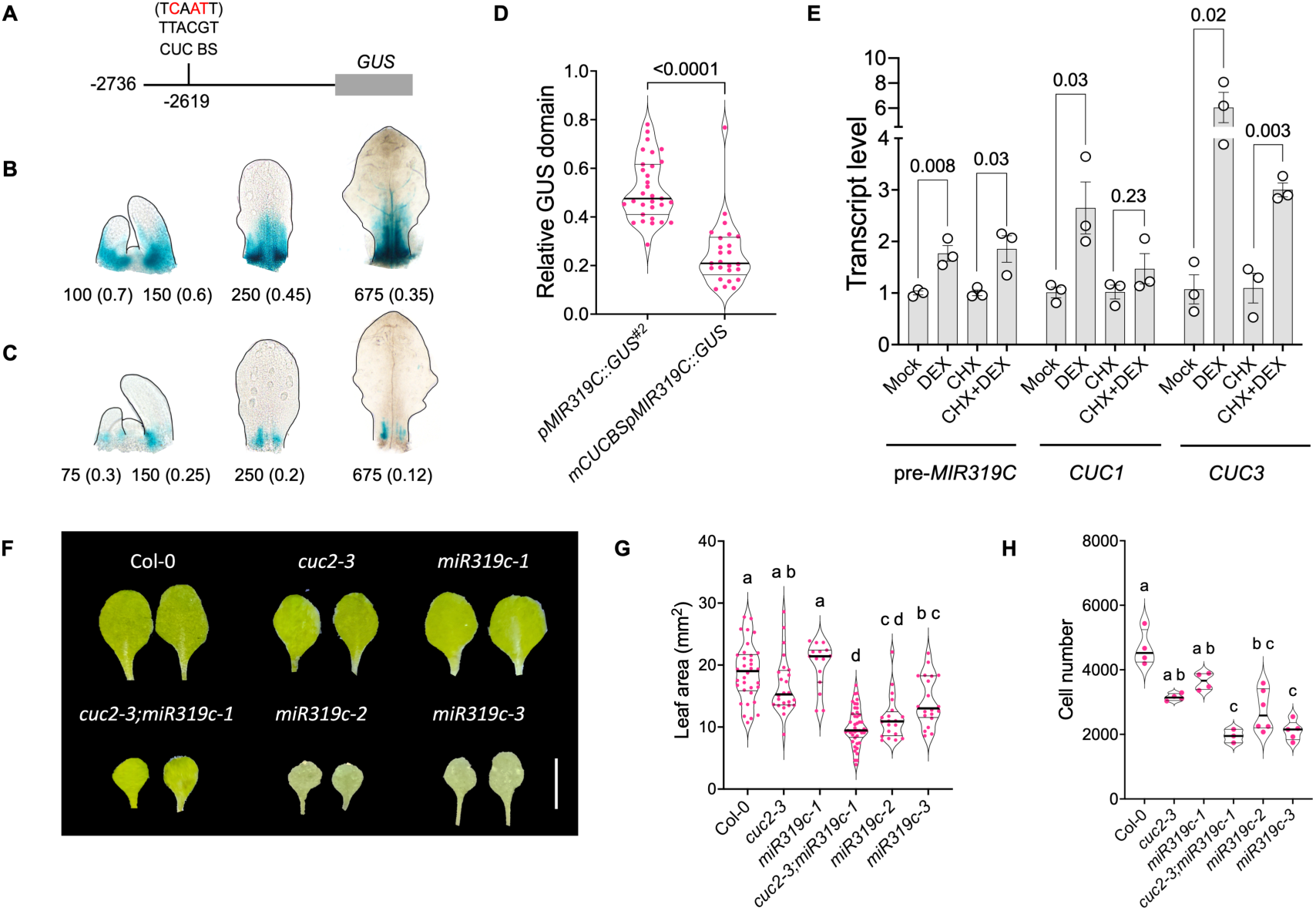
Role of *CUC2*-*MIR319C* module in leaf cell proliferation. **(A)** Schematic representation of the *pMIR319C*::*GUS* construct with mutated bases (red) on the putative CUC-binding site (TTACGT). Numbers indicate nucleotide positions upstream to transcription start site. The *GUS* coding sequence is indicated by a grey box. **(B-C)** Bright field images of processed shoot apices (left) and leaf primordia expressing *GUS-*based reporter transgene under wildtype (*pMIR319C::GUS*, **B**) or mutated (*mCUCBSpMIR319C::GUS*, **C**) *MIR319C* promoters. Numbers below the images indicate leaf length (larger leaf primordia are considered for the apices); relative GUS domain values are indicated within parentheses. Black lines highlight the boundaries. **(D)** Violin plot representations of relative GUS domain in the 50-750 μm long leaf primordia (N, 25-31) on 4^th^-8^th^ nodes of indicated genotypes. Differences between samples are indicated by *p*-value above the data points; unpaired *t-*test with Welch’s correction for differences in variance was used. **(E)** Transcript level of genes indicated below the X-axis labels in 10-day old *pRPS5A::CUC2-GR-HA* seedlings treated for 2 hours with ethanol (Mock) or 12 µM dexamethasone (DEX) and/or 10 µM cycloheximide (CHX). Error bars represent SEM of three biological replicates. Pairwise comparisons were performed using unpaired *t-*test; *p*-values are shown above the data bars. **(F)** Representative images of mature first pair of leaves from 25-day old plants of indicated genotypes. Scale bar, 5 mm. **(G)** Areas (Y-axis) of mature first pair of leaves of indicated genotypes. N, 15-44. Significant differences among the samples are indicated by lower-case letters. p<0.0001. One-way ANOVA, followed by Dunn’s multiple comparison test were performed to determine the significant differences among samples. **(H)** Total number of pavement cells in mature first pair of leaves of the indicated genotypes. Cell number was determined by dividing leaf area by cell area (100-120 cells per leaf) averaged over 3-6 leaves per genotype. Significant differences among samples are indicated by lower-case letters. p<0.01; one-way ANOVA, followed by Tukey’s multiple comparison test were performed.

To further test whether CUC2 directly activates *MIR319C* transcription, we induced CUC2 activity in 10-day-old *pRPS5A::CUC2-GR-HA* seedlings (Tian et al. 2014) by treating them with dexamethasone (DEX) for two hours in the presence of the protein synthesis inhibitor cycloheximide (CHX) and measured pre-*MIR319C* level by RT-qPCR. CUC2 induction increased the transcript of *CUC3* (**Fig. 2E**), one of its known direct targets (Maugarny-Calès et al. 2019), but not of *CUC1*, in the presence of CHX. Similar to positive control, CUC2 induction also elevated the pre-*MIR319C* transcript level by ∼2-fold in the presence of CHX, suggesting that no new protein synthesis is required for activating *MIR319C* transcription by CUC2. These results indicate that CUC2 binds to the *MIR319C* locus and activates its transcription in leaf primordia.

### *CUC2* promotes leaf cell proliferation and leaf size in miR319-dependent manner

The mature miR319 produced by the *MIR319A*, *MIR319B* & *MIR319C* genes promotes cell proliferation and leaf size by degrading the transcripts of five *JAW-TCP*s (Palatnik et al. 2003; Efroni et al. 2008; Koyama et al. 2010; Challa et al. 2019). If CUC2 promotes miR319 accumulation, inactivation of either *CUC2* or *MIR319C* is expected to yield smaller leaves with fewer cells. To test this, we analysed mature leaves of the *CUC2* loss-of-function allele *cuc2-3* (Nikovics et al. 2006), and a *MIR319C* loss-of-function allele available in the Arabidopsis stock centre (https://arabidopsis.info/; established as a line and named here as *miR319c-1*) (**Fig. S3A**). While *miR319c-1* mutant did not show any noticeable reduction in mature leaf area (**Fig. 2F, 2G)** or epidermal cell area (**Fig. S4A, S4B**), mature leaf size of *cuc2-3* was reduced by ∼20% compared to Col-0 (**Fig. 2F, 2G)**. The weak phenotypes of single mutants are perhaps because of the hypomorphic nature of the mutant alleles or due to the presence of their redundant partners (Nag et al. 2009; Hasson et al. 2011). To test the former possibility for *MIR319C*, we generated novel knockout mutants of *MIR319C* using CRISPR zCas9i editing approach (See methods; Grützner et al. 2021, Stuttmann et al. 2021), and recovered the *miR319c-2* allele with complete, and the *miR319c-3* allele with partial deletion of the region that produces mature miR319c (**Fig. S5**). Both these alleles produced mature leaves with nearly 40% smaller area compared to wild type (**Fig. 2F, 2G)** without a noticeable change in average cell area (**Fig. S4A, S4B)**, resulting in reduced cell number in the mutants (**Fig. 2H**). These results show that miR319c activity is required for leaf cell proliferation.

To explore the genetic relationship between *CUC2* and its target *MIR319C* in regulating cell proliferation, we generated the double mutant line *cuc2-3;miR319c-1* by crossing weak alleles of these two genes and analysed its leaf size. Mature *cuc2-3;miR319c-1* leaves were nearly half the size of their respective parents (**Fig. 2F, 2G**), though the average area of their pavement cells remained unaltered (**Fig. S4A, S4B**). The *cuc2-3;miR319c-1* leaf blade was made up of ∼2000 pavement cells whereas the Col-0 leaves had ∼4500 cells (**Fig. 2H**), demonstrating that smaller area of the double mutant leaves is due to reduced cell proliferation. The corresponding cell numbers for the *cuc2-3* and *miR319c-1* leaves also reduced compared to the wildtype (∼3100 for *cuc2-3* and ∼3500 for *miR319c-1*), but to a much less extent (**Fig. 2H**). In combination with the biochemical and expression analysis described above, we interpreted results of these genetic analysis as CUC2 promoting cell proliferation in leaf primordia in a miR319c-dependent manner.

During leaf morphogenesis, cell proliferation commences throughout the incipient primordia, followed by concurrent proliferation and expansion in distinct zones, with all cells finally ceasing to proliferate and undergoing secondary expansion (Donnelly et al. 1999; Andriankaja et al. 2012; Fox et al. 2018). To determine the stage at which cell proliferation in the *cuc2-3*;*miR319c-1* leaves diverges from wild type, we performed confocal imaging of propidium iodide-stained leaf primordia at 1, 3, and 5 days after initiation (DAI) and analysed their cellular parameters using MorphographX software (Barbier de Reuille et al. 2015; Strauss et al. 2022). The basipetal growth pattern of *cuc2-3*;*miR319c-1* leaves were comparable with that of Col-0 at all the early growth stages studied (**Fig. 3A**). Further, the total number and average size of the pavement cells also remained unaltered in the mutant leaf primordia (**Fig. S6A-D**), suggesting that loss of *CUC2* or *MIR319C* does not affect cell proliferation or its spatial pattern at early growth stage.

**Figure 3:**
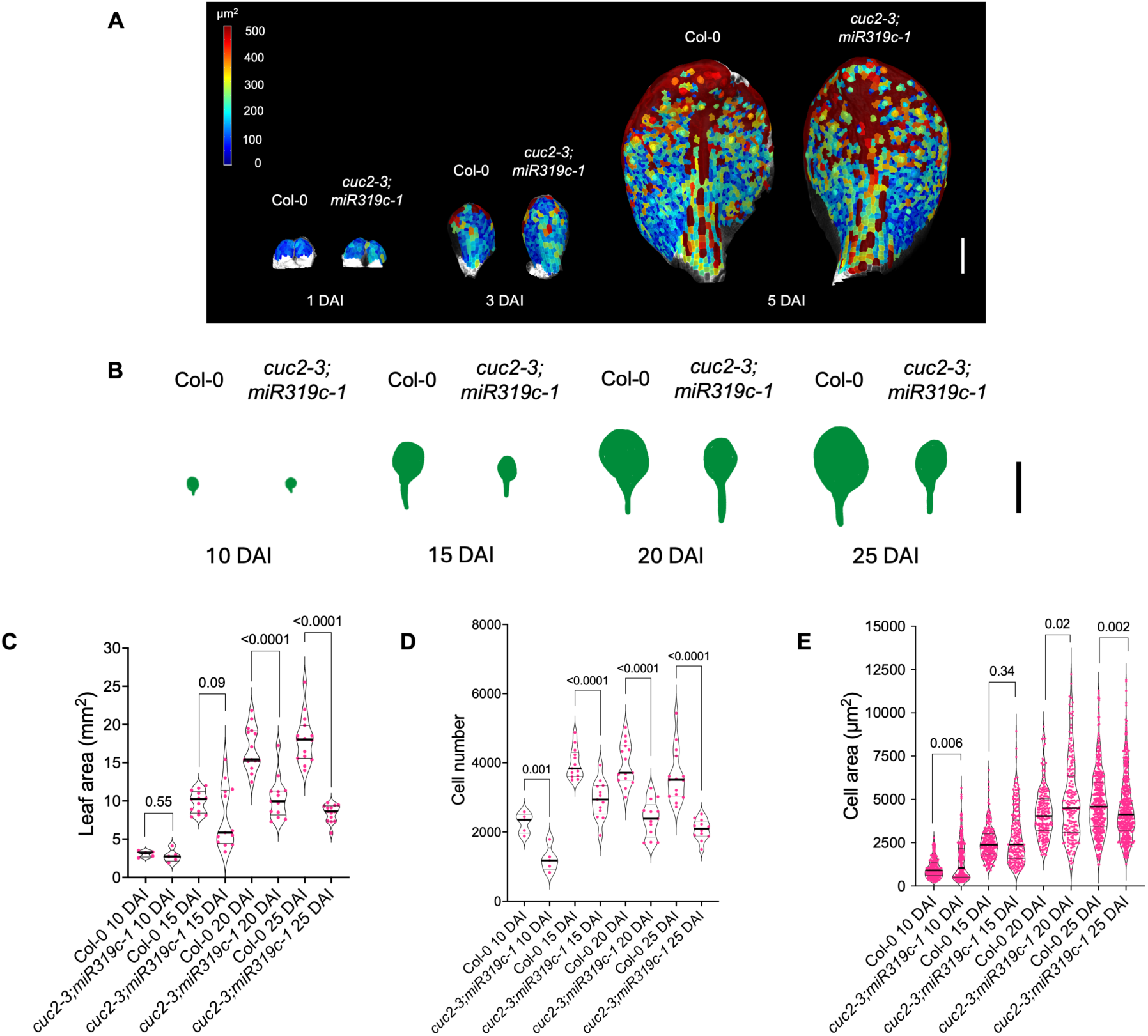
Growth kinematics of *cuc2-3;miR319c-1* leaves. **(A)** Heat maps of cell areas in the first pair of Col-0 and *cuc2-3;miR319c-1* leaves at 1, 3, and 5 days after initiation (DAI). Scale bar, 100 μm. **(B)** Impressions of Col-0 and *cuc2-3;miR19c-1* leaves at indicated DAI. Scale bar, 5 mm. **(C-E)** Violin plot representations of leaf area (**C**), total pavement cell number (**D**), and average pavement cell area (**E**) of mature first leaf pair of Col-0 and *cuc2-3;miR319c-1* leaves at DAI indicated on the X-axes. Bold horizontal lines within the datapoints represent median. Differences among samples are indicated by *p*-values above data points; unpaired *t-*test with Welch’s correction for differences in variance (**C, D**) or Mann Whitney U test (**E**) were used. N, 5-15 leaves (**C, D**) or 199-447 cells (**E**).

To examine whether the *CUC2*-*MIR319C* module promotes cell proliferation at later stages, we analysed leaf and cell growth kinematics by measuring leaf area, pavement cell area, and total cell number of Col-0 and *cuc2-3;miR319c-1* leaves at 10, 15, 20, and 25 DAI. The analysis revealed that average leaf area of *cuc2-3;miR319c-1* was reduced to nearly half at later stages of growth, i.e., 15 DAI onwards (**Fig. 3B, 3C**), with concomitant and corresponding decrease in total cell number at all stages (**Fig. 3D**). The average area of the *cuc2-3;miR319c-1* epidermal cells, though increased with time, remained comparable to that of Col-0 (**Fig. 3E**), suggesting that the reduction of leaf size in the mutant was primarily due to reduced cell proliferation. Though average cell size remained same, the mutant leaves had higher proportion of larger cells at all growth stages measured (**Fig. S7**), pointing at precocious differentiation. Together, these results demonstrate that the *CUC2-MIR319C* module promote cell proliferation and delays differentiation in leaf primordia.

*MIR319A* and *MIR319B*, the two homologues of *MIR319C*, are required for leaf serration and loss of function of both the genes produces smooth margin (Koyama et al. 2017). To examine whether *MIR319A* and *MIR319B* also regulate cell proliferation in leaf primordia along with *MIR319C*, we generated higher order mutants of these three genes by crossing *miR319a;miR319b* double mutant with *miR319c-1* and determined their leaf area and pavement cell parameters. Mature *miR319a;miR319b;miR319c-1/+* and *miR319a;miR319b/+;miR319c-1* mutant plants produced leaves that are less than one-third the size of the leaves of their respective parents and of Col-0, while the *miR319a;miR319b;miR319c-1* triple homozygous leaves were ∼4.5 times smaller than Col-0 leaves (**Fig. 4A, 4C**). Reduction in cell number alone accounted for smaller leaf area (**Fig. 4D**), with the average size of the pavement cells remaining nearly unaltered (**Fig. S8A, S8B**). Together, these results suggest a functional redundancy among three *MIR319* genes in promoting leaf cell proliferation.

**Figure 4:**
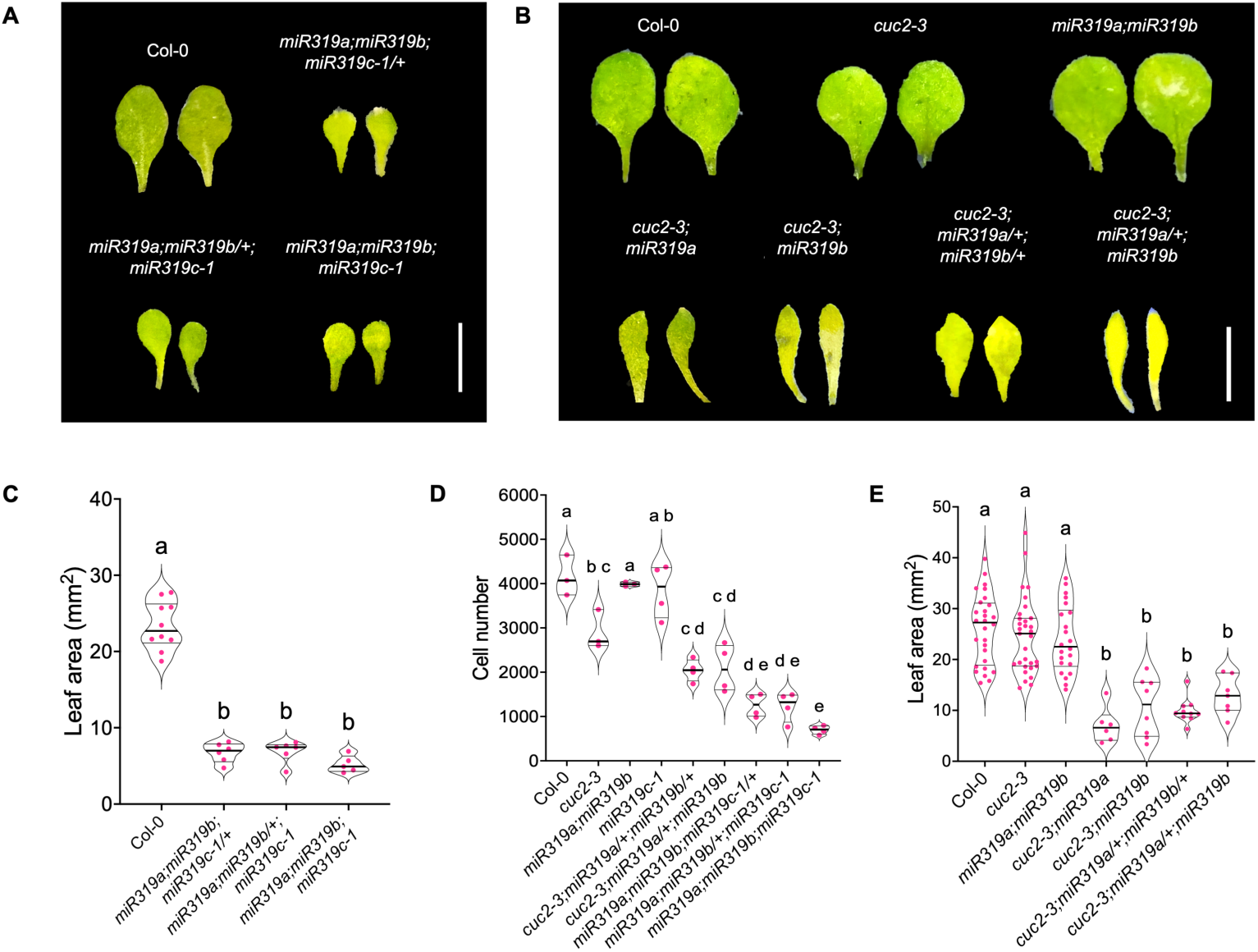
Genetic interaction between *CUC2* and *MIR319* genes in regulating leaf cell proliferation. (A, **B)** Representative images of mature first pair of leaves dissected from 25-day old plants of the indicated genotypes. Scale bar, 5 mm. **(C)** Areas of mature first pair of leaves of the indicated genotypes. N, 5-10 leaves. Significant differences among the samples are indicated by lower-case letters. p<0.0001; one-way ANOVA, followed by Dunnett’s T3 multiple comparison test were performed to determine the significant differences among samples. **(D)** Total number of pavement cells in mature first pair of leaves of the indicated genotypes, as derived from dividing leaf area by cell area (75-100 cells per leaf) averaged from 3-4 leaves per genotype. Significant differences among the samples are indicated by lower-case letters. p<0.05; one-way ANOVA, followed by Tukey’s multiple comparison test was performed to determine the significant differences among samples. **(E)** Areas of mature first pair of leaves of the indicated genotypes. N, 6-31 leaves. Significant differences among the samples are indicated by lower-case letters. p<0.01; one-way ANOVA, followed by Dunnett’s T3 multiple comparison test was performed to determine the significant differences among samples.

To further investigate a possible genetic interaction between *MIR319A/ MIR319B* and *CUC2* in regulating leaf cell proliferation, we combined *miR319a;miR319b* genotype with *cuc2-3* and *cuc2-1* and studied the leaf phenotype in higher-order mutants. Whereas most (∼96%) F2 individuals of the *miR319a;miR319b* x *cuc2-3* genetic cross displayed parental phenotype, a small fraction of them (32 out of 821) showed aberrant phenotypic features including smaller leaves and fused cotyledons (**Fig. S9A**), the latter being characteristic to *cuc* loss of function mutants (Aida et al. 1997; Hibara et al. 2006). Similar phenotypic aberrations of varying proportions were also noticed among the F2 individuals of the *miR319a;miR319b* x *cuc2-1* cross (**Fig. S9A**). More importantly, nearly one third individuals of the genetically established *cuc2-3;miR319a* & *cuc2-3;miR319b* homozygous populations, and of *cuc2-3;miR319a/+;miR319b/+* & *cuc2-3;miR319a/+;miR319b* populations also showed smaller leaves (2-4 folds) and fused cotyledons (**Fig. 4E**, **Fig. S9B**), suggesting a genetic interaction among these genes. Reduced leaf area in *cuc2-3;miR319a/+;miR319b/+* & *cuc2-3;miR319a/+;miR319b* lines was accompanied with >2-fold decrease in the number of pavement cells (**Fig. 4D**) without any increase in their size (**Figs. S8A, S8B**). Thus, the *CUC2-MIR319* pathway promotes cell proliferation in early leaf primordia.

### *CUC-MIR319C-JAW-TCP* module regulates leaf size

Since mature miR319 suppresses the function of five *JAW-TCP* genes by degrading their transcripts, CUC2-mediated *MIR319* activation and promotion of cell proliferation is expected to be dependent of JAW-TCP activity. We tested this by carrying out genetic interaction studies between loss and gain-of-function lines of *CUC2* and *JAW-TCP*s. The gain-of-function line of *CUC2*, *cuc2-1D*, forms larger leaves with increased cell number (Larue et al. 2009), perhaps due to enhanced *MIR319C* level (**Fig. 1D-F**) and a consequential decrease in *JAW-TCP* level. Activation of an miR319-resistant, dominant form of TCP4 within its endogenous expression domain in the *Col-0;GR^#1^* x Col-0 F1 plants reduces leaf size by limiting cell proliferation (**Fig. 5A, 5C**, Challa et al. 2016). TCP4 activation in the *Col-0;GR^#1^* x *cuc2-1D* F1 plants reduced the size of the bigger *cuc2-1D* leaves to the same level as in the Col-0 background (**Fig. 5A, 5C**), thus suggesting that *JAW-TCP*s function downstream to *CUC2*. Overexpression of miR319 under a CUC2-insentitive promoter in the *pBLS>>MIR319A* leaves expectedly led to larger lamina (**Fig. 5B, 5D;** Efroni et al. 2008; Challa et al. 2019). However, additionally mutating *CUC2* in this genetic background did not further alter the leaf size (**Fig. 5B, 5D**). Taken together, these results support the conclusion that *CUC2* promotes leaf size by reducing *JAW-TCP* level in a miR319-dependent manner.

**Figure 5:**
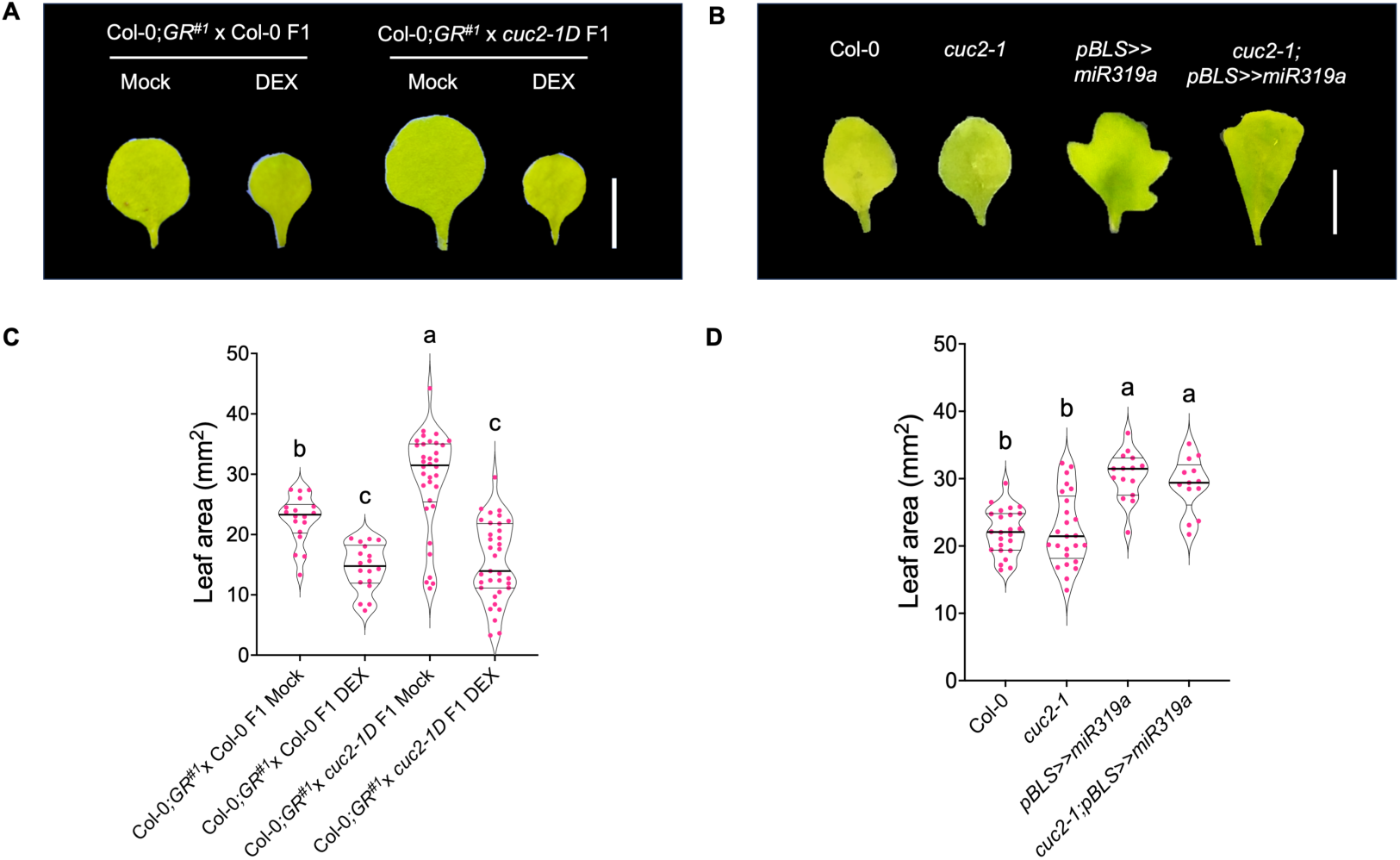
Genetic interaction between *CUC2* and *JAW-TCP* genes in regulating leaf size. (A, **B)** Representative images of mature first leaves from 25-day old plants of the indicated genotypes. Scale bar, 5 mm. **(C, D)** Areas of mature first pair of leaves of indicated genotypes. N, 13-35. Significant differences among samples are indicated by different lower-case letters. *p*<0.01; two-way ANOVA, followed by Tukey’s post hoc test (**C**) or Dunnett’s T3 multiple comparison test (**D**) were performed to determine the significant differences among samples. Mock and DEX (**A**, **C**) indicate continuous treatment with solvent (Mock) or 12 μM dexamethasone (DEX).

### *CUC2* and *JAW-TCP* activities together establish *MIR319C* expression domain in early leaf primordia

Members of the *JAW-TCP* clade establish early expression patterns of *MIR319C* by suppressing its transcription in the distal incipient leaf primordia, and of the *CUC* genes by restricting them to the proximal domain of the early shoot lateral organs (Koyama et al. 2007; Challa & Rath et al. 2021; Shankar et al. 2023). To examine whether JAW-TCPs modulate *CUC2* transcription in establishing the endogenous *MIR319C* expression pattern during leaf growth, we investigated *pCUC2::GUS* activity (Nikovics et al. 2006) in *jaw-D*, a knockdown line of *JAW-TCP*s (Palatnik et al. 2003), and found that the *CUC2* expression domain remained unaltered in the *jaw-D* leaf primordia (**Fig. 6A**). Further, activation of a dominant TCP4 by dexamethasone treatment in the Col-0 x Col-0;*GR^#1^* F1 seedlings did not change the *pCUC2::GUS* domain either (**Fig. S8C**), suggesting that JAW-TCPs do not affect *CUC2* transcription during the growth of leaf primordia.

**Figure 6:**
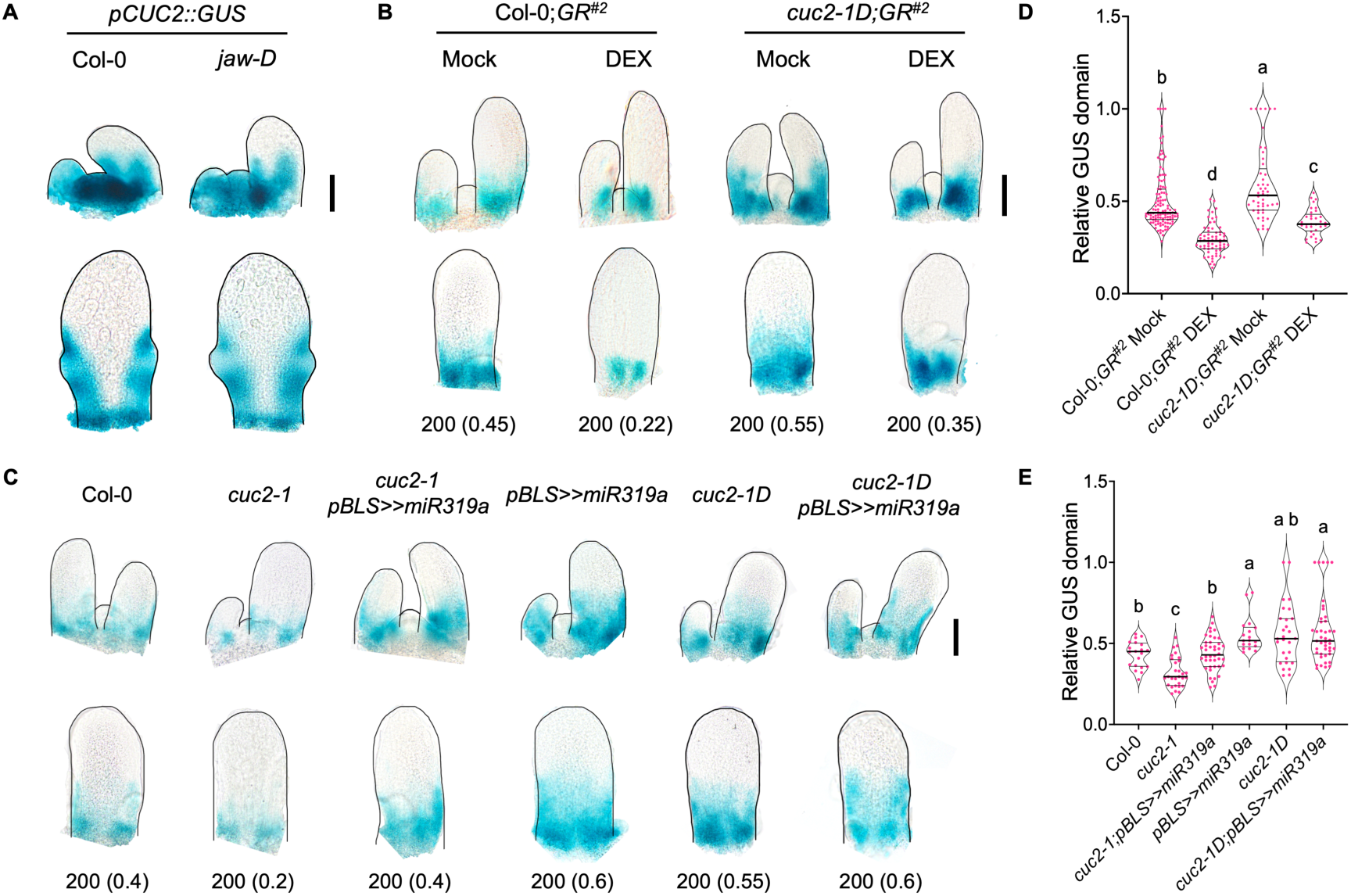
Regulation of *MIR319C* expression by *CUC2* and *JAW-TCP* genes in leaf primordia. **(A)** Bright field images of cleared shoot apices (top panel) and 6^th^ leaf primordia (bottom panel) from 10-day old Col-0 and *jaw-D* seedlings expressing *pCUC2::GUS* transgene. Black lines highlight borders of the images. Scale bar, 50 μm. **(B, C)** Bright-field images of cleared shoot apices (top panel) and 6^th^ leaf primordia (bottom panel) dissected from 10-day old seedlings of indicated genotypes expressing the *pMIR319C::GUS* transgene. Black lines highlight borders. Scale bar, 50 μm. Numbers below the images represent leaf lengths in μm and relative GUS domain values (in parentheses). Mock and DEX in (**B, D**) indicate continuous treatment with solvent (Mock) or 12 μM dexamethasone (DEX). **(D, E)** Relative *pMIR319C::GUS* domain values in 50-350 μm leaf primordia plotted for the indicated genotypes. Differences among samples are indicated by lower-case letters. p<0.05. N varied from 33-114 (**D**) or 18-43 (**E**). Two-way ANOVA followed by Tukey’s post hoc test (**D**), or one-way ANOVA followed by Dunnett’s post hoc test (**E**) were performed to determine the significant differences among samples.

To further investigate how JAW-TCPs and CUC2 together establish *MIR319C* expression pattern, we determined *MIR319C* promoter activity in leaf primordia where the level of both *CUC2* and *JAW-TCP*s were altered. As was reported earlier, activation of a dominant TCP4 protein in the Col-0;*GR^#2^*leaves, where an miR319-resistant TCP4 is induced by dexamethasone treatment in the wildtype background (Shankar et al. 2023), *pMIR319C::GUS* domain was reduced and restricted more proximally (**Fig. 6B, 6D**). However, the extent of restriction was less when TCP4 was induced in the *cuc2-1D;GR^#2^* leaf primordia, indicating that the endogenous *MIR319C* expression domain is established by independent and opposing activities *CUC2* and *JAW-TCPs*. In support of this, downregulation of the *JAW-TCP*s rescued the reduced *MIR319C* expression of the *cuc2-1* leaves (**Fig. 6C, 6E**). However, downregulation of *JAW-TCP*s failed to further increase the already extended *MIR319C* domain in *cuc2-1D* (**Fig. 6C, 6E**), perhaps suggesting that either *CUC2* activation or *JAW-TCP* inactivation results in the maximum extent of *MIR319C* domain at the leaf growth stages studied here.

## Discussion

Regulation of organ size is a fundamental question in developmental biology. In angiosperm species such as Arabidopsis, leaves grow initially via cell proliferation, followed by a combination of proliferation and expansion that are spatially separated along the length of the primordia. Finally, all pavement cells attain their maximum and fairly uniform size throughout the leaf lamina in a mature leaf. While the timing of the transition from cell proliferation to differentiation has been shown to influence final cell number of leaf size, its molecular basis remains somewhat unclear. Here, we report that the *CUC2-MIR319-JAW-TCP* module plays a key role in promoting pavement cell proliferation by delaying its transition to differentiation, thereby impacting leaf size. Our data is consistent with the model (**Fig. 7A**) where CUC2 activates *MIR319C* transcription by binding to a cis element in the *MIR319C* upstream regulatory region (**Figs. 1 and 2**), leading to an elevated level of miR319c activity in the incipient leaf primordia. As the primordia grow, *CUC2* expression becomes progressively restricted towards leaf-SAM boundary where it continues to activate *MIR319C* expression. The miR319c activity at the leaf base prevents JAW-TCP accumulation and sustains cell proliferation. JAW-TCPs first accumulate in the distal cells of the growing primordia due to a yet unknown mechanism, and feedback to repress *MIR319C* transcription within their expression domain (Shankar et al. 2023). The opposing activities of CUC2 and JAW-TCPs *MIR319C* transcription independently establish the endogenous miR319 domain in early leaf primordia (**Fig. 6**). Further experiments are needed to test whether CUC2 directly binds to this element in planta or if additional factors are required for CUC2 binding on the *MIR319C* locus.

**Figure 7:**
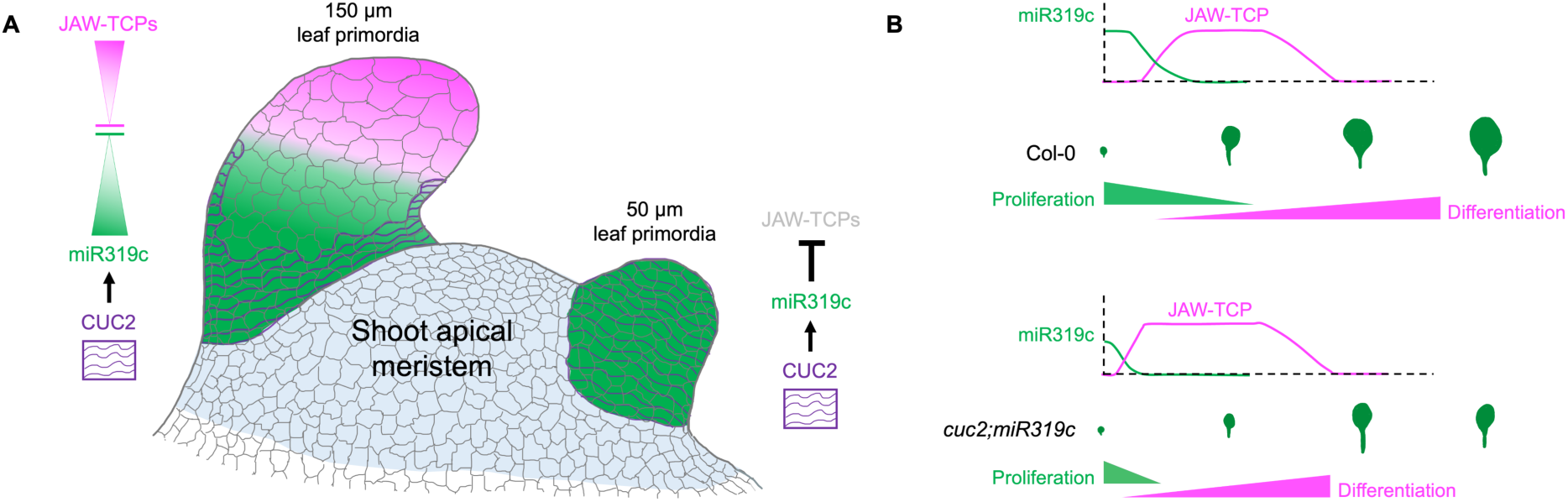
Schematic representation of proposed model for CUC2-miR319c mediated upregulation of cell proliferation in leaf primordia. **(A)** CUC2 (purple wavy lines) expression throughout the incipient leaf primordium (on the right) where it directly activates *MIR319C* expression (green). The mature miR319c ensures cell proliferation by preventing accumulation of *JAW-TCP* transcripts (bold black T-line). As the primordium grows out of the shoot apical meristem (SAM), CUC2 expression gets restricted towards the SAM-leaf boundary (primordium on the left), where it maintains miR319c activity that continues to repress *JAW-TCP*s, thus establishing the basal cell proliferation domain (left panel). At this stage, JAW-TCP proteins accumulate at the distal region (magenta on the left panel), and directly repress *MIR319C* transcription (inverted magenta triangle) (Shankar et al. 2023), thus promoting proliferation-to-differentiation transition in distal cells by distinct mechanisms (Challa et al. 2019). Thus, the *MIR319C* expression domain is established via the antagonistic regulation by JAW-TCP and CUC2 proteins. Image credits: Hand-drawn outline of the shoot apex adapted from Du et al. 2018 (https://doi.org/10.1016/j.molp.2018.06.006) **(B)** In Col-0 (upper panel), high miR319c level (green line) inhibit JAW-TCP (magenta line), ensuring proliferation (green triangle) at the base of early leaf primordia. As primordia develop, miR319c domain declines, whereas JAW-TCP domain increases concurrently, committing proliferating cells to differentiation (magenta triangle). JAW-TCP activity remains high in the growing leaf, promoting increased cell differentiation and expansion-driven growth. As leaves approach maturity, JAW-TCP level decline, and all cells complete differentiation and growth by expansion. In *cuc2;miR319c* leaf primordia (bottom panel), reduced miR319c levels lead to an early rise in JAW-TCP activity, causing a premature transition from proliferation to differentiation and expansion-driven growth. This reduction in proliferation potential decreases the final cell number, ultimately resulting in a smaller mature leaf with fewer cells.

### Role of the *CUC2-MIR319C* module in spatiotemporal regulation of leaf cell proliferation

In our study, loss of *CUC2* results in only a modest reduction in leaf size, primarily due to a decrease in cell number (**Fig. 2**). This may in part be attributed to the partial downregulation of miR319c, as evidenced by reduced promoter activity in the *cuc2* loss-of-function mutant (**Fig. 1**). The strong reduction in cell number observed in the *cuc2-3;miR319c-1*, and in the *miR319c-2* or *miR319c-3* null mutants, support this hypothesis (**Fig. 2**). We speculate that the *cuc2-3;miR319c-1* mutant has lower levels of miR319c than the individual parental alleles that are perhaps hypomorphic. Since *MIR319C* expression is restricted to the basal half of leaf primordia by 3 DAI (∼250 μm long primordia; **Fig. S1**), and JAW-TCPs are active at the distal end by this stage (Shankar et al. 2023), we investigated if the cell proliferation pattern is altered in *cuc2-3;miR319c-1* during early leaf development. However, no significant changes in cell number, cell areas or cell size distribution were observed up to 5 DAI (∼500 μm long primordia; **Fig. 3**). Difference in leaf area and cell number started appearing only by 10 DAI onwards (**Fig. 3**), a stage when *CUC2* and *MIR319C* expressions are restricted to the base of the petiole (**Fig. S1**). These observations suggest that the *CUC2-MIR319-JAW-TCP* module does not influence cell division rate or the establishment of the initial leaf growth pattern but rather affects the duration of pavement cell proliferation by delaying the transition.

JAW-TCPs are known to promote the proliferation-to-differentiation transition via auxin dependent and independent genetic pathways (Challa et al. 2019). The reduction in miR319 level in *cuc2-3;miR319c-1* leaves is likely to enhance JAW-TCP level, leading to premature differentiation of cells that would otherwise continue to proliferate in the wild-type primordia. Over time, this effect compounds, ultimately leading to a detectable reduction in cell number at later stages, leading to smaller leaves. This is further supported by our kinematic analysis, which revealed that *cuc2-3;miR319c-1* leaves consistently have fewer cells at 10 DAI and at subsequent stages of growth (**Fig. 3**), indicating an early transition from proliferation to differentiation. This interpretation corresponds with the increased proportion of larger pavement cells in the mutant leaves as early as at 10 DAI (**Fig. S7**). Additional evidence comes from the genetic interactions between the mutants of *CUC2* and *JAW-TCPs*, where ectopic JAW-TCP activity reduced the size of *CUC2*-overexpressing leaves, and miR319 overexpression rescued smaller-size phenotype of the *cuc2-1* leaves (**Fig. 5**). Overall, our findings support the model (**Fig. 7B**) where CUC2 activation of *MIR319C* maintains the proliferation potential of leaf cells from the base to the transition zone in growing leaves by downregulating JAW-TCP activity. In the absence of this regulation, the proliferation phase is shortened, resulting in fewer cells and smaller leaves in *cuc2-3;miR319c-1*.

### Redundancy among *MIR319* genes in regulating leaf size

Individual role of the three *MIR319* genes in regulating leaf cell proliferation were not clear until now due to functional redundancy among them and due to the availability of null mutants. Defects in leaf area and cell number in *miR319c-2; miR319c-3* plants (**Fig. 2**) revealed a key role for all the three *MIR319C* genes in promoting cell proliferation via downregulation of JAW-TCP level. Reduced leaf area and cell number in the triple *miR319a;miR319b;miR319c-1* mutant but not in the *miR319a;miR319b* mutant or in *miR319c-1* mutant (**Fig. 4**) suggest that *MIR319A* and *MIR319B* also play a role in maintaining cell proliferation. Moreover, *MIR319A/ MIR319B* displayed a strong genetic interaction with *CUC2* in promoting cell number and leaf size (**Fig. 4**). This is likely due to the additional reduction in the mature miR319 pool in *cuc2-3;miR319a;miR319b* leaves resulting from the combined loss of all three *MIR319* genes. It is also possible that CUC2 activates *MIR319A/ MIR319B* genes in leaf primordia, in addition to *MIR319C*. Interestingly, the *cuc2-3;miR319a; miR319b* mutants exhibited reduced cell area and, in some cases, distinctly altered pavement cell morphology (**Fig. S8**), suggesting varying degrees of contribution from *MIR319A/ MIR319B* genes to leaf growth. This variation may result from spatiotemporal differences in *MIR319* expression during leaf development. In future, detailed analysis of *MIR319A/ MIR319B* expression pattern (Nag et al. 2009) combined with growth kinematic study of their mutant leaves, will help clarify these possibilities.

### Context-dependent role of *CUC2* in cell proliferation

*CUC2* perhaps plays a context-dependent role in regulating cell proliferation and establishing growth polarity during organ morphogenesis. During embryogenesis , *CUC2* suppresses cell proliferation at the boundary between SAM and emerging cotyledon primordia to ensure cotyledon separation (Aida et al. 1997; Ishida et al. 2000). Similarly, *CUC2* also inhibits cell proliferation in at the sinus region of leaf primordia and thereby promotes serration development (Nikovics et al. 2006; Maugarny-Calès et al. 2019; Hasson et al. 2011), a boundary function parallel to its effect on cotyledon separation during embryogenesis. Here, we demonstrate that *CUC2* rather promotes cell proliferation in the medial region of the lamina base through the *miR319-JAW-TCP* pathway. This dual role of *CUC2* as both a suppressor and a promoter of cell proliferation could be influenced by developmental context and tissue-specific factors. Additionally, our results support the idea that *CUC2* functions as a growth polarity organizer (Kuchen et al. 2012; Sarvepalli et al. 2019), where it drives differential growth between the proximal and the distal regions of the leaf primordia via *MIR319C* activation. This also raises an interesting possibility that tissue boundaries can serve as growth organizers during organ morphogenesis in plants (Bhatia et al. 2023; Hu et al. 2024). Future research could explore mechanisms that link boundary gene network and growth regulatory modules to test this possibility.

## Methods

### In silico screening for SAM and leaf primordia-specific transcription factors

To identify SAM and leaf primordia-specific TFs, we took advantage of studies that generated cellular-resolution maps of genes whose mRNA is actively translated in different domains within the SAM and leaf primordia (Tian et al. 2014; 2019). We considered genes that are active in the *CLV3*, *WUS*, *UFO* expression domains of the SAM, *LAS* expression domain of the SAM-leaf boundary, and the *AS1* expression domain of the leaf primordia. We overlapped the 4426 SAM and leaf primordia-specific genes with the list of 1956 genome-wide Arabidopsis TFs (Pruneda-Paz et al. 2014) and obtained 343 TFs expressed explicitly in the SAM and leaf primordia domains (**Suppl Data Set. 3**). We also scanned the 2.7 kb URR of *MIR319C* for TF binding sites using the publicly available transcription factor binding site (TFBS) prediction tool (PlantPAN 3.0 -http://plantpan.itps.ncku.edu.tw). We recovered several potential DNA-binding motifs on the *MIR319C* promoter, including TCP, KNOX, YABBY, NAC and HD-like TF-binding consensus. We listed 49 representatives from the top 5 TF families recovered from the TFBS prediction software (**Suppl Data Set. 4**). Next, we combined this list with the previously obtained 343 SAM and leaf primordia-specific TFs to get a final list of 378 TF candidates for the yeast one-hybrid screen (**Suppl Data Set. 1)**.

### Yeast transformation

Yeasts were transformed with the desired DNA constructs (plasmids) by the high-efficiency LiAc-PEG method described by Walhout and Vidal 2001, with minor modifications. A single colony of host strain was inoculated in 10 ml YPAD (Bacto yeast extract-1g, Tryptone-1g, Dextrose-2g, and Adenine sulphate-10mg per 100ml) and incubated overnight at 30°C with constant shaking. The next day, the suitable culture volume was inoculated into 200 ml of fresh YPAD solution to an initial OD of 0.2 and kept at 30°C with constant shaking until an OD of 0.6-0.8 was reached. Cells were pelleted down at 700 g for 10 min and washed twice with 5 ml of sterile water. The cells were washed with 2.5 ml of LiAc-TE (0.1 M LiAc, Tris-10 mM, EDTA-1mM) solution. Cells were finally resuspended in 2 ml of LiAc-TE solution. A mixture containing 280 μL of 50% PEG (Mol wt 3350, Sigma-Aldrich, USA), 36 μL of 1 M LiAc in TE, 6 μL of 10 mg/ml boiled carrier DNA (D-3159, Sigma-Aldrich, USA) was prepared and added to 1.5 ml microfuge tubes containing 60 μL cells and 100 ng of DNA construct. The cell suspension was vortexed briefly and incubated at 30°C with constant shaking for 30 min. Cells were transferred into a water bath maintained at 42°C for 30 min. Cells were pelleted, washed twice with sterile water, resuspended in 100 μL sterile water, and plated onto appropriate media.

For the Y1H screen, transformations were carried out in 96 well-format described by Walhout and Vidal, 2001. Yeast cells were prepared as described in the high-efficiency protocol. 20 μL of cell suspension (containing LiAc-PEG-carrier DNA) were transferred to each well of a sterile U-bottom 96-well plate (3795, Corning, USA). 100 ng of plasmid and 100 μL of TE/LiAc/PEG solution (50% PEG and 0.1 M LiAc in TE) were individually added and resuspended carefully using a multichannel pipette. 96-well plate was incubated at 30°C with constant shaking for 30 min. Cells were transferred to a water bath maintained at 42°C for 30 min. Cells were pelleted at 700 g for 5 min. To each well, 120 μL of sterile water was added, and 100 μL was removed directly to get rid of PEG in the supernatant. Finally, cells were resuspended in the remaining 20 μL water and spotted on appropriate media.

### Yeast bait, prey, and diploid generation

The full length (2.7 kb) and the non-overlapping promoter fragments, i.e., -2694-1866 (*p*^distal^), -1866-1107 (*p*^middle^), and -1107+35 (*p*^proximal^) of *MIR319C* were amplified from *pMIR319C-P4P1R* entry clone using Taq DNA polymerase and cloned into pENTR/D-TOPO vector (K240020, Invitrogen, USA) according to the manufacturer’s instructions. The entry clones were confirmed by Sanger sequencing before cloning upstream to the *minimalPro::HIS3* cassette of *pMW#2* (Gift from Ram Yadav, IISER Mohali, INDIA) by LR reaction. Y1H baits were generated according to the protocol described by Fuxman Bass et al. 2016. Briefly, the *pMIR319C* containing *pMW#2* reporter constructs were linearized with *Xho*I (R0146, New England Biolabs, USA) and integrated into the mutant *HIS3* locus of the *YM4271* host strain (Gift from Ram Yadav, IISER Mohali, INDIA) using the high-efficiency LiAc-PEG transformation method described by Walhout and Vidal, 2001. Following transformation, yeast cells were plated on the synthetic complete media lacking histidine (Sc-His) media (Walhout and Vidal, 2001) and incubated at 30°C till colonies appeared. Post growth, genomic DNA was extracted individually from 3-4 colonies, and the integration of the *MIR319C* promoter fragment was confirmed by PCR. All primer sequences used for cloning are mentioned in **Suppl Data Set. 5**.

To generate the prey strains for the Y1H screen, each of the 384 TF-encoding plasmids was individually purified from corresponding glycerol stocks of the 1956 DEST-U Arabidopsis TF ORFeome collection (CD4-89, ABRC, USA) using QIAprep Spin Miniprep Kit. 100 ng of purified plasmid were individually transformed into Yalpha1867 strain (Gift from Dr. Ram Yadav, IISER Mohali, INDIA) using the high throughput 96-well format transformations protocol described earlier. Following transformation, the cells were spotted on synthetic complete media lacking tryptophan (Sc-Trp) media and incubated at 30°C till colonies appeared.

To generate the diploids used in the Y1H assay, prey TF-encoding plasmids was individually purified from corresponding glycerol stocks of the 1956 DEST-U Arabidopsis TF ORFeome collection (CD4-89, ABRC, USA) using QIAprep Spin Miniprep Kit. 100 ng of purified plasmid were individually transformed into *pMIR319C* baits using the high-efficiency LiAc-PEG method described earlier. Following transformation, the cells were spotted on synthetic complete media lacking histidine and tryptophan (Sc-His-Trp) media and incubated at 30°C till colonies appeared.

### Yeast one-hybrid (Y1H) screen/assay

The auto activity of yeast baits on Sc-His due to minimal *HIS3* expression was determined by spotting them on Sc-His media containing 0-30 mM 3-Amino-1,2,4-triazole (3-AT) (A8056, Sigma-Aldrich, USA). The minimum concentration of 3-AT that completely inhibits the growth of the bait was considered for the Y1H screen/ assay. For the mating-based Y1H screen, the diploids generated by mating were resuspended in sterile 96-well plates containing Sc-Trp-His media and allowed to grow in shaking incubators for 24 hrs at 30°C. After 24 hrs of growth, OD was measured. Volumes corresponding to equal OD were spotted on Sc-Trp-His+3AT media, incubated at 30°C, and scored for growth on days 3-5 post-incubation. Spots containing at least 5 colonies were considered positive for growth. For the Y1H screen, spots showing growth on all three concentrations of 3-AT, i.e., 20, 25, and 30 mM by day 5, or growth on 20 and 25 mM 3-AT by day 3 were considered positive for binding. For the co-transformation-based Y1H assay, yeast baits were transformed with TF encoding plasmids and selected on Sc-Trp-His media. Following selection, the cells were resuspended in Sc-Trp-His media and allowed to grow in shaking incubators for 24 hrs at 30°C. After 24 hrs of growth, OD was measured. Volumes corresponding to equal OD were spotted on Sc-Trp-His media with respective concentrations of 3-AT and incubated at 30°C.The plates were inspected for growth and imaged over days 3-5 post-incubation.

### Plant materials

*Arabidopsis thaliana* ecotype Columbia-0 (Col-0) was used as the wild-type control. All the lines used in this study are in Col-0 genetic background. The genotypes used in this study are described in **Suppl Data Set. 6**. The genotypes *cuc2-1* and *pBLS>>miR319a* (Originally in L*er*; Efroni et al. 2008; Hasson et al. 2011) was brought into the Col-0 background by back crossing three times. *pMIR319C::GUS^#1^* (Nag et al. 2009) was established in various mutant backgrounds by crossing and was confirmed by GUS assays in the F3/F4 generation. *cuc2-3;miR319c-1*, *cuc2-3;miR319ab* and *miR319a;miR319b;miR319c-1* mutants were generated by crossing respective mutant lines *cuc2-3* (Nikovics et al. 2006), *miR319ab* (Koyama et al. 2017), *miR319c-1* (SAIL_647_B03/ CS827925) and confirmed by genotyping in F3 generation. *cuc2-1D;GR^#2^* was established by crossing *cuc2-1D* (Larue et al. 2009) and Col-0;*GR^#2^* (Shankar et al. 2023) mutants and confirmed by phenotyping in F3 generation. All genotyping primers are listed in **Suppl Data Set. 5**.

### Plant growth condition and chemical treatments

For GUS assays and phenotyping experiments, seeds were surface-sterilized with 0.05% SDS containing 70% ethanol solution for 10 min, followed by wash with 100% ethanol twice and sown on 1X Murashige and Skoog (MS), 0.8% agar medium (PT011, HiMedia Laboratories, India) containing 0.025% ethanol (Mock) or 6 μM dexamethasone (DEX) (D4902, Sigma-Aldrich, USA). After sowing, seeds were kept for stratification for three days in the dark at 4°C and then shifted to a plant growth chamber (I-41LL, Percival Scientific, USA) set at 22°C, 60% relative humidity, long-day conditions (16h/8h light/dark period, 100 μmol.m^-2^.s^-1^). 14-day old seedlings were transplanted onto a soil mixture containing Soilrite and Perlite (Keltech Energies, India) in a 3:1 ratio. Plants were grown till maturity in growth rooms with the conditions as described for growth chamber.

For transient DEX and cycloheximide (CHX) treatments, 9 day-old mock (1X MS, 0.8% agar medium containing 0.025% ethanol) grown seedlings were transferred onto petri dishes filled with 0.5X MS medium, supplemented with 1% sucrose, 1X Gamborg B5 vitamins (PL018, HiMedia Laboratories, India), 12 μM DEX, and/or 10 μM CHX (C7698, Sigma-Aldrich, USA), incubated for 2 hours and harvested under liquid N2 prior to RNA isolation.

For confocal imaging experiments, seeds were stratified in the dark for two days at 4°C to synchronize germination. Plants were grown on petri dishes filled with 0.5X MS medium, supplemented with 0.8% agar, 1% sucrose, and 0.1% Plant Protective Mixture (Plant Cell Technology) in a growth chamber under long-day conditions (16h/8h light/dark period, 95 µmol.m^-2^.s^-1^) at 22 ± 1°C with 60-70% relative humidity.

### DNA constructs and plant transformation

For the generation of *pMIR319C* deletion and *mCUCBSpMIR319C::GUS* constructs, a 2.7 kb genomic fragment corresponding to the *MIR319C* URR from the start of pre-*MIR319C* was amplified from Col-0 genomic DNA using Phusion High Fidelity DNA polymerase (M0530, New England Biolabs, USA) and cloned into modified *P4P1R* entry vector (a kind gift from Ram Yadav, IISER Mohali, India) by gateway BP reaction (11789020, Invitrogen, USA) to generate the *pMIR319C-P4P1R* construct. To introduce mutation at the CUC-binding site, we utilized a PCR-based site directed mutagenesis approach. Briefly, forward, and reverse primers carrying mutations in CUC-binding site (mCUCBS) was first used in a PCR reaction with *pMIR319C-P4P1R* template using Phusion DNA polymerase. Post PCR reaction, the product was digested with *Dpn*I (R0176, New England Biolabs, USA) and transformed into *E. coli* and selected on LB agar containing 100 mg/l ampicillin. Plasmids were purified from resulting colonies using QIAprep Spin Miniprep Kit (Qiagen, Germany) and mCUCBS was confirmed by Sanger sequencing. The entry clone *mCUCBSpMIR319C-P4P1R* was recombined into *R4L1pGWB532* destination vector (Gift from Tsuyoshi Nakagawa, Shimane University, Japan) by LR reaction (11791020, Invitrogen, USA) to generate the *mCUCBSpMIR319C::GUS* construct. Sequence information of primers used to generate the constructs are provided in the **Suppl Data Set. 5**. *miR319c* targeting CRISPR constructs were generated based on zCas9i modular cloning approach (Grützner et al. 2021, Stuttmann et al. 2021). The zCas9i cloning kit: MoClo-compatible CRISPR/Cas9 cloning kit with intronized Cas9 was a gift from Sylvestre Marillonnet and Johannes Stuttmann (Addgene kit #1000000171). The guide RNAs-sgRNA1 (5’-AGTGACTGTGTGAATGATG-3’) and sgRNA2 (5’-TATCTGTGTTTGGACTGAA-3’) were cloned into pDGE332 and pDGE334 shuttle vectors and assembled into pDGE347 vector according to cloning manual v5 of the zCas9i cloning kit.

*mCUCBSpMIR319C::GUS* and the *miR319c* CRISPR constructs were integrated into Col-0 background by the *Agrobacterium tumefaciens*-mediated floral dip method (Clough and Bent, 1998). For *mCUCBSpMIR319C::GUS* line generation, seeds were harvested and screened for the ability to grow on MS media in presence of 20 mg/l hygromycin. At least five independent lines were established for each construct and confirmed by genotyping in T3 generation. Sequence information of primers used for genotyping are provided in the **Suppl Data Set. 5**. For generating CRISPR knockouts of *miR319c*, seeds were harvested and screened for red fluorescence and for the ability to grow on MS media in presence of 10 mg/l BASTA. T2 seedlings from ten independent T1 plants were screened for deletions in miR319c region by Sanger sequencing. Two independent lines, *miR319c-2* and *miR319c-3*, carrying deletions in the miR319c region were established in T3 generation after counter selecting for lack of red fluorescence in T2 population.

### β-glucuronidase (GUS) assay

GUS assays were performed as described earlier (Shankar et al. 2023). Briefly, samples were collected in ice-cold 90% acetone, incubated at room temperature for 20 min, followed by washing with ice-cold staining buffer (50 mM sodium phosphate, pH 7.0, 0.2% Triton X-100, 5 mM potassium ferrocyanide, and 5 mM potassium ferricyanide). Fresh staining buffer was added with 2 mM X-Gluc (R0851, Thermo Scientific, USA) and incubated overnight at 37°C in dark. Staining buffer was replaced with 70% ethanol, and 70% ethanol was changed 3-4 times in regular intervals to clear the tissue. Shoot apices and leaf primordia were dissected and mounted on a glass slide containing lactic acid under the microscope and were imaged using Olympus BX51 trinocular DIC microscope fitted with ProgResC3 camera and ProgResCapture Pro2.6 software (Jenoptik, Germany) with the following settings, exposure-65ms, brightness-10 units, contrast-35 units, colour temperature-5000K, image resolution-(1040 x 770 2X Bin), while keeping all other settings in default mode.

### GUS domain, leaf area and pavement cell area quantification

The extent of reporter gene expression in the *pMIR319C::GUS* and *mCUCBSpMIR319C::GUS* transgenic lines were determined using a pixel density quantification method reported in Shankar et al., 2023. Images of 6 and 9-day old *cuc2-3;miR319ab* seedlings were captured using LEICA EZ4 E stereomicroscope equipped with LAS EZ software. Areas of the dissected mature first pair of leaves from 25-day old plants were determined using the polygon selection tool of the ImageJ software. For pavement cell area measurements, dissected mature first pair of leaves were first incubated in 7:1 absolute ethanol: glacial acetic acid clearing solution. After 24 hours, clearing solution was replaced with 1M KOH solution for 30 minutes, washed twice with sterile water and incubated in 37% D-lactic acid solution until tissue became completely transparent. The clarified leaf samples were mounted on a glass slide using D-lactic acid and imaged at 20x magnification on a Olympus BX51 trinocular DIC microscope fitted with ProgResC3 camera and ProgResCapture Pro2.6 software (Jenoptik, Germany). Cell areas were determined using the polygon selection tool of the ImageJ software. The average cell number was derived using the leaf area and cell area of leaves taken for the experiment. Silhouettes of cleared leaves were used for representative purposes using an exact scale of the image captured.

### Confocal microscopy and imaging

For imaging of the first pair of leaf, one cotyledon was removed using fine tweezers or an injection needle from two-day-old plants to expose the initiating leaf primordium for imaging of leaf at 1 day after initiation (DAI). The seedlings were stained with 1 mg/mL Propidium Iodide (PI) solution for 2-3 minutes and then were rinsed with water. The dissected plants then were placed horizontally in Ø60 mm Petri dishes filled with 0.5X MS medium, supplemented with 1.5% agar, 1% sucrose, and 0.1% Plant Protective Mixture (Plant Cell Technology). Plants were immersed in water containing 0.1% PPM for imaging. Three to four replicates were imaged using the PI staining and at least half of the abaxial leaf surface was imaged at 1 DAI, 3 DAI and 5 DAI respectively.

All confocal imaging was performed with an LSM980 upright confocal microscope using a long-distance water-dipping objective (AP 20X/1.0; Zeiss) at the confocal facility of MechanoDevo team (RDP, ENS de Lyon). Excitation was performed using a diode laser with 561 nm for PI. Images were collected at 580-650 nm for PI with a GaAsP detector and a beam splitter operating at 488 and 561 nm. Confocal stacks were acquired at 512×512 resolution and 16-bit image depth, with 1.4 µm distance in Z-dimension. For samples larger than the field of view, multiple overlapping stacks were obtained and later stitched together using MorphoGraphX (Barbier de Reuille et al., 2015; Strauss et al., 2022).

### Confocal Image analysis

Cellular growth and expression quantifications were conducted using MorphoGraphX. Stacks were processed as described previously (Le Gloanec et al. 2022). After digitally removing the trichomes when necessary, the organ surface was detected with the ‘Edge detect’ tool with a threshold of 20000 with ‘closing’ to fill holes, followed by the ‘Edge detect angle’ (10000). An initial 5 µm cube size mesh was then created and subdivided three times before projecting the membrane signal (2-5 µm). Cell segmentation was performed manually. Metrics such as area cell size were computed as described before (Barbier de Reuille et al. 2015; Sapala et al. 2018).

### RNA isolation and RT-qPCR

Total RNA was extracted using the TRIzol method described earlier (Challa et al. 2016). 10 μg of RNA was used for DNase (EN0521, Thermo Scientific, USA) treatment. 1.5 μg DNase-treated RNA was converted to cDNA using RevertAid Reverse Transcriptase (EP0441, Thermo Scientific, USA) and 2.5 μM oligo dT according to the manufacturer’s instructions. Quantitative PCR reactions with 25 ng cDNA and 0.4 μM primers were set up using the DyNAmo Flash SYBR Green RT-qPCR kit (F415L, Thermo Scientific, USA), according to the manufacturer’s instructions. 10 μl qPCR reactions were carried out in QuantStudio™ 6 Flex Real-Time PCR System, 384-well format (Applied Biosystems, USA). Results were analyzed using the inbuilt QuantStudio6 Pro software, and ΔΔCT values were determined after normalization with *PROTEIN PHOSPHATASE 2A SUBUNIT A3 (PP2AA3*). Relative fold changes in transcript levels were calculated using the formula 2^-ΔΔCT^. All primer sequences are provided in **Suppl Data Set. 5**.

### Statistical analysis

All graphical representations and statistical analysis were performed using GraphPad Prism 9 software (GraphPad, USA). First, samples were tested for Gaussian distribution using the Shapiro-Wilk normality test. If samples were normally distributed, pairwise comparisons were performed using unpaired *t-*test. Welch’s correction was applied if variances were different among the samples. Statistical differences among multiple genotypes were determined by ordinary One-Way ANOVA followed by Tukey’s multiple comparisons test or One-Way ANOVA followed by Dunnet’s T3 post-hoc test, depending on whether variances were significantly different among the genotypes. If the samples failed the normality test, then pairwise comparisons were carried out by Mann Whitney test. Statistical differences among multiple genotypes were determined by One-Way ANOVA followed by Dunn’s post-hoc test. To compare statistical differences among multiple genotypes subjected to ethanol (Mock) or dexamethasone (DEX) treatments, we used Two-Way ANOVA followed by Tukey’s post-hoc test. *p-*values obtained from the respective tests are indicated above the data bars in several plots. For plots with a large number of comparisons, statistical differences among samples are indicated by different letters. Raw data and statistics summary of all the plots are provided in **Suppl Data Set. 7**.

### List of Supplementary Data Files

Suppl Data Set 1. List of 378 candidate TFs used in yeast one hybrid screen (TF mini library)

Suppl Data Set 2. List of 57 TF positives from yeast one hybrid screen

Suppl Data Set 3. List of SAM and leaf primordia-specific TFs (343)

Suppl Data Set 4. List of TFs from TFBS analysis on *MIR319C* promoter (49)

Suppl Data Set 5. List of primers used in this study (5’ to 3’)

Suppl Data Set 6. List of genotypes used in this study

Suppl Data Set 7. Raw data and statistics summary

### Accession numbers

TAIR (The Arabidopsis Information Resource, www.arabidopsis.org) accession numbers of the significant genes used in this study are *AT4G23713* (*MIR319A*), *AT5G41663* (*MIR319B*), *AT2G40805* (*MIR319C*), *AT3G15170* (*CUC1*), *AT5G53950* (*CUC2*), *AT1G76420* (*CUC3*), *AT3G15030* (*TCP4*).

## Acknowledgements

We sincerely thank Ram Yadav (IISER Mohali, India) for vectors and strains used in Y1H assays, Koyama Tomotsugu (Bioorganic Research Institute, Kyoto, Japan) for the *miR319ab* seeds, Patrick Laufs (Institut Jean Pierre Bourgin, INRA Versailles, France) for the *pCUC2::GUS* line, Yuling Jiao (Peking University, Beijing, China) for *pRPS5A::CUC2-GR-HA* line. We thank Sachin Kotak, Yadukrishnan Premachandran and Anurag Sharma of IISc Bangalore, India, for helpful discussions during the preparation of this manuscript.

## Author contributions

N.S. initiated the project, designed & performed most experiments, analysed & interpreted the results, organized the figures, wrote the first draft of the manuscript and contributed to its finalization. V.M. performed confocal microscopy and growth kinematics experiments, collected data and assisted in organizing the figures. A.G. assisted in some of the RT-qPCR experiments and imaged mature leaves, young seedlings for some of the phenotyping experiments. V.M. and A.G. performed most of the cell area measurements. S.R. generated *miR319c-2,* and *miR319c-3* lines, performed some of the GUS expression experiments. A.J.P. established the *miR319c-1* line. A.V. made CRISPR constructs for generation of *miR319c-2* and *miR319c-3* lines. O.H. contributed to data interpretation and conclusions, corrected the manuscript and provided the infrastructure for confocal imaging and analysis. U.N contributed to designing experiments and data interpretation, guided the first six co-authors, corrected and finalised the manuscript.

## References

1. Aida M, Ishida T, Fukaki H, Fujisawa H, Tasaka M Genes involved in organ separation in Arabidopsis: An analysis of the cup-shaped cotyledon mutant. Plant Cell 1997: 9: 841–857

2. Aida M, Ishida T, Tasaka M Shoot apical meristem and cotyledon formation during Arabidopsis embryogenesis: interaction among the CUP-SHAPED COTYLEDON and SHOOT MERISTEMLESS genes. Development. 1999:126: 1563–1570

3. Andriankaja M, Dhondt S, DeBodt S, Vanhaeren H, Coppens F, DeMilde L, Mühlenbock P, Skirycz A, Gonzalez N, Beemster GTS, et al Exit from Proliferation during Leaf Development in Arabidopsis thaliana: A Not-So-Gradual Process. Dev Cell. 2012:22: 64–78

4. Bar M, Ori N Leaf development and morphogenesis. Development. 2014: 141: 4219–4230

5. Barbier de Reuille P., Routier-Kierzkowska A.L., Kierzkowski D., Bassel GW., Schüpbach T., Tauriello G., Bajpai N, Strauss S, Weber A, Kiss A et al. MorphoGraphX: A platform for quantifying morphogenesis in 4D. Elife. 2015: 4: 05864. doi:10.7554/eLife.05864

6. Barton MK Twenty years on: The inner workings of the shoot apical meristem, a developmental dynamo. Dev Biol. 2010: 341: 95–113

7. Bhatia N, Wilson-Sánchez D, Strauss S, Vuolo F, Pieper B, Hu Z, Rambaud-Lavigne L, and Tsiantis M. Interspersed expression of CUP-SHAPED COTYLEDON2 and REDUCED COMPLEXITY shapes Cardamine hirsuta complex leaf form. Current Biology. 2023:33(14):2977–2987.e6. 10.1016/j.cub.2023.06.037

8. Bilsborough GD, Runions A, Barkoulas M, Jenkins HW, Hasson A, Galinha C, Laufs P, Hay A, Prusinkiewicz P, Tsiantis M. Model for the regulation of Arabidopsis thaliana leaf margin development. Proc Natl Acad Sci U S A. 2011: 108: 3424–3429

9. Challa KR, Aggarwal P, Nath U. Activation of *YUCCA5* by the Transcription Factor TCP4 Integrates Developmental and Environmental Signals to Promote Hypocotyl Elongation in Arabidopsis. Plant Cell. 2016: 28: 2117–2130

10. Challa KR, Rath M, Nath U. The CIN-TCP transcription factors promote commitment to differentiation in Arabidopsis leaf pavement cells via both auxin-dependent and independent pathways. PLoS Genet. 2019: 15: 1–30

11. Clough SJ, Bent AF. Floral dip: A simplified method for Agrobacterium-mediated transformation of Arabidopsis thaliana. Plant J. 1998: 16: 735–43

12. Donnelly PM, Bonetta D, Tsukaya H, Dengler RE, Dengler NG. Cell cycling and cell enlargement in developing leaves of Arabidopsis. Dev Biol. 1999: 215: 407–419

13. Du F, Guan C, Jiao Y. Molecular Mechanisms of Leaf Morphogenesis. Mol Plant. 2018: 11: 1117–1134

14. Efroni I, Blum E, Goldshmidt A, Eshed Y. A Protracted and Dynamic Maturation Schedule Underlies Arabidopsis Leaf Development. the Plant Cell Online. 2008: 20: 2293–2306

15. Fox S, Southam P, Pantin F, Kennaway R, Robinson S, Castorina G, Sánchez-Corrales YE, Sablowski R, Chan J, Grieneisen V, et al. Spatiotemporal coordination of cell division and growth during organ morphogenesis. PLoS Biol. 2018: 16: e2005952

16. Fuxman Bass JI, Pons C, Kozlowski L, Reece-Hoyes JS, Shrestha S, Holdorf AD, et al. A gene-centered C. elegans protein–DNA interaction network provides a framework for functional predictions. Mol Syst Biol. 2016: 12: 884

17. Gázquez A, Beemster GTS. What determines organ size differences between species? A meta-analysis of the cellular basis. New Phytologist. 2017: 215: 299–308

18. Gonzalez N, Vanhaeren H, Inzé D. Leaf size control: Complex coordination of cell division and expansion. Trends Plant Sci. 2012: 17: 332–340

19. Grutzner R, Martin P, Horn C, Mortensen S, Cram EJ, Lee-Parsons CWT, Stuttmann J, Marillonnet S. High-efficiency genome editing in plants mediated by a Cas9 gene containing multiple introns. Plant Commun. 2021:2:100135.

20. Hasson A, Plessis A, Blein T, Adroher B, Grigg S, Tsiantis M, Boudaoud A, Damerval C, Laufs P. Evolution and diverse roles of the CUP-SHAPED COTYLEDON genes in Arabidopsis leaf development. Plant Cell. 2011: 23: 54–68

21. Hepworth J, Lenhard M. Regulation of plant lateral-organ growth by modulating cell number and size. Curr Opin Plant Biol. 2014: 17: 36–42

22. Hibara KI, Karim MR, Takada S, Taoka KI, Furutani M, Aida M, Tasaka M. Arabidopsis CUP-SHAPED COTYLEDON3 regulates postembryonic shoot meristem and organ boundary formation. Plant Cell. 2006:18: 2946–2957

23. Hu ZL, Wilson-Sánchez D, Bhatia N, Rast-Somssich MI, Wu A, Vlad D, McGuire L, Nikolov LA, Laufs P, Gan X, et al. A CUC1/auxin genetic module links cell polarity to patterned tissue growth and leaf shape diversity in crucifer plants. Proc Natl Acad Sci U S A. 2024:121(26). 10.1073/pnas.2321877121

24. Ishida T, Aida M, Takada S, Tasaka M. Involvement of CUP-SHAPED COTYLEDON genes in gynoecium and ovule development in Arabidopsis thaliana. Plant Cell Physiol. 2000: 41: 60–67

25. Kalve S, De Vos D, Beemster GTS. Leaf development: a cellular perspective. Front Plant Sci . 2014:5: 1–25

26. Koyama T, Furutani M, Tasaka M, Ohme-Takagi M.TCP Transcription Factors Control the Morphology of Shoot Lateral Organs via Negative Regulation of the Expression of Boundary-Specific Genes in Arabidopsis. Plant Cell. 2007: 19: 473–484

27. Koyama T, Mitsuda N, Seki M, Shinozaki K, Ohme-Takagi M. TCP transcription factors regulate the activities of ASYMMETRIC LEAVES1 and miR164, as well as the auxin response, during differentiation of leaves in Arabidopsis. Plant Cell. 2010: 22: 3574–3588

28. Koyama T, Sato F, Ohme-Takagi M. Roles of miR319 and TCP transcription factors in leaf development. Plant Physiol. 2017: 175: 874–885

29. Kuchen EE, Fox S, De Reuille PB, Kennaway R, Bensmihen S, Avondo J, et al. Generation of Leaf Shape Through Early Patterns of Growth and Tissue Polarity. 2012; 335:1092–6

30. Larue CT, Wen J, Walker JC. A microRNA-transcription factor module regulates lateral organ size and patterning in Arabidopsis. Plant Journal. 2009: 58: 450–463

31. Le Gloanec C, Collet L, Silveira SR, Wang B, Routier-Kierzkowska AL, and Kierzkowski D. Cell type-specific dynamics underlie cellular growth variability in plants. Development. 2022: 149: dev200783.

32. Lindemose S, Jensen MK, De Velde J Van, O’Shea C, Heyndrickx KS, Workman CT, Vandepoele K, Skriver K, Masi F De. A DNA-binding-site landscape and regulatory network analysis for NAC transcription factors in Arabidopsis thaliana. Nucleic Acids Res. 2014: 42: 7681–7693

33. Maugarny-Calès A, Cortizo M, Adroher B, Borrega N, Gonçalves B, Brunoud G, Vernoux T, Arnaud N, Laufs P. Dissecting the pathways coordinating patterning and growth by plant boundary domains. PLoS Genet. 2019: 15: 1–30

34. Maugarny-Calès A, Laufs P. Getting leaves into shape: A molecular, cellular, environmental and evolutionary view. Development. 2018: 145: dev161646

35. Nag A, King S, Jack T. miR319a targeting of TCP4 is critical for petal growth and development in Arabidopsis. Proceedings of the National Academy of Sciences. 2009: 106: 22534–22539

36. Nikovics K, Blein T, Peaucelle A, Ishida T, Morin H, Aida M, Laufs P. The balance between the MIR164A and CUC2 genes controls leaf margin serration in Arabidopsis. Plant Cell. 2006: 18: 2929–2945

37. Ooka H, Satoh K, Doi K, Nagata T, Otomo Y, Murakami K, Matsubara K, Osato N, Kawai J, Carninci P, et al. Comprehensive Analysis of NAC Family Genes in Oryza sativa and Arabidopsis thaliana. DNA Research. 2003: 10: 239–247

38. Palatnik JF, Allen E, Wu X, Schommer C, Schwab R, Carrington JC, Weigel D. Control of leaf morphotgenesis by microRNAs. Nature. 2003: 425: 257–263

39. Pruneda-Paz, JL, Breton, G, Nagel, DH, Kang, SE, Bonaldi, K, Doherty, CJ, Ravelo, S, Galli, M, Ecker, JR, and Kay, SA. A Genome-Scale Resource for the Functional Characterization of Arabidopsis Transcription Factors. Cell Rep. 2014: 8: 622–632

40. Rodriguez RE, Mecchia MA, Debernardi JM, Schommer C, Weigel D, Palatnik JF. Control of cell proliferation in Arabidopsis thaliana by microRNA miR396. Development. 2010: 137:103–12

41. Rodriguez RE, Debernardi JM, Palatnik JF. Morphogenesis of simple leaves: Regulation of leaf size and shape. Wiley Interdiscip Rev Dev Biol. 2014: 3: 41–57

42. Rubio-Somoza I, Zhou CM, Confraria A, Martinho C, Von Born P, Baena-Gonzalez E, Wang JW, Weigel D. Temporal control of leaf complexity by miRNA-regulated licensing of protein complexes. Current Biology. 2014: 24: 2714–2719

43. Sapala, A, Runions A, Routier-Kierzkowska AL, Gupta MD, Hong L, Hofhuis H, Verger S, Mosca G, Li CB, Hay A, et al. Why plants make puzzle cells, and how their shape emerges. Elife. 2018: 7: e32794.

44. Sarvepalli K, Das Gupta M, Challa KR, Nath U. Molecular cartography of leaf developmentC—Crole of transcription factors. Curr Opin Plant Biol. 2019: 47: 22–31

45. Sarvepalli K, Nath U. CIN-TCP transcription factors: Transiting cell proliferation in plants. IUBMB Life. 2018: 70: 718–731

46. Schneider M, Van Bel M, Inzé D, Baekelandt A. Leaf growth – complex regulation of a seemingly simple process. Plant Journal. 2023: 117: 1018–1051

47. Scofield S, Murison A, Jones A, Fozard J, Aida M, Band LR, Bennett M, Murray JAH. Coordination of meristem and boundary functions by transcription factors in the SHOOT MERISTEMLESS regulatory network. Development. 2018: 145: dev157081

48. Shankar N, Sunkara P, Nath U. A double-negative feedback loop between miR319c and JAW-TCPs establishes growth pattern in incipient leaf primordia in Arabidopsis thaliana. PLoS Genet. 2023: 19: 1–25

49. Strauss S, Runions A, Lane B, Eschweiler D, Bajpai N, Trozzi N, Routier-Kierzkowska AL, Yoshida S, Rodrigues da Silveira S, Vijayan A, et al. Using positional information to provide context for biological image analysis with MorphoGraphX 2.0. Elife. 2022: 11: e72601

50. Stuttmann J, Barthel K, Martin P, Ordon J, Erickson, J L, Herr R, Ferik F, Kretschmer C, Berner T, Keilwagen J. et al. Highly efficient multiplex editing: one-shot generation of 8× Nicotiana benthamiana and 12× Arabidopsis mutants. Plant J. 2021: 106, 8–22.

51. Tian C, Zhang X, He J, Yu H, Wang Y, Shi B, Han Y, Wang G, Feng X, Zhang C, et al. An organ boundaryCenriched gene regulatory network uncovers regulatory hierarchies underlying axillary meristem initiation. Mol Syst Biol. 2014: 10: 755

52. Tian, C., Wang, Y., Yu, H., He, J., Wang, J., Shi, B., Du, Q., Provart, N.J., Meyerowitz, E.M., and Jiao, Y. A gene expression map of shoot domains reveals regulatory mechanisms. Nat. Commun. 2019: 10: 1–12.

53. Vroemen CW, Mordhorst AP, Albrecht C, Kwaaitaal MACJ, De Vries SC. The CUP-SHAPED COTYLEDON3 gene is required for boundary and shoot meristem formation in Arabidopsis. Plant Cell. 2003: 15: 1563–1577

54. Wang Q, Hasson A, Rossmann S, Theres K. Divide et impera: Boundaries shape the plant body and initiate new meristems. New Phytologist. 2016: 209: 485–498

