## Supplementary Figures for "*MIR319C* activation by CUP-SHAPED COTYLEDON2 delays cell proliferation-to-differentiation transition in Arabidopsis leaf primordia"

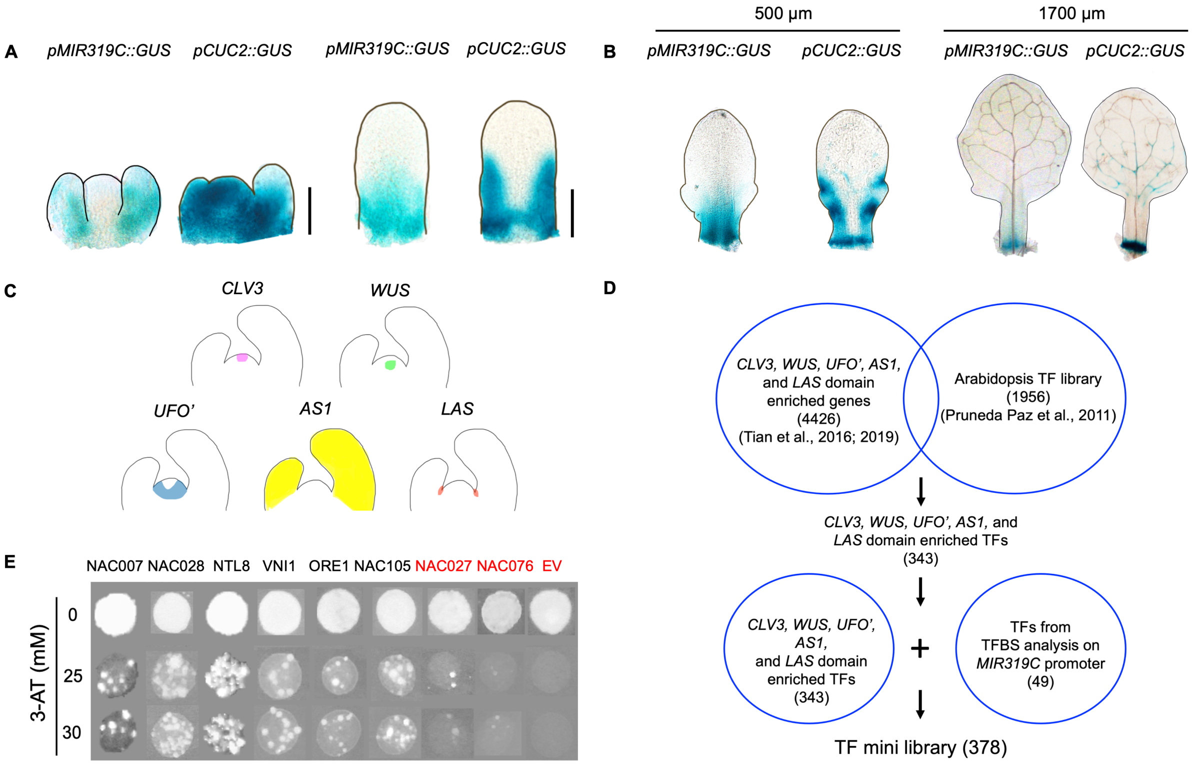


**Figure S1 (Supports Fig. 1): Yeast one-hybrid screen of TFs with *MIR319C* upstream regulatory region as bait. (A)** Bright field images of 10-day old, cleared shoot apices (left panel) and 6$^{th}$ leaf primordia (right panel) expressing *GUS*-based reporter transgene for *MIR319C* or *CUC2* promoters as indicated. Black outlines of the SAM and leaf primordia are hand drawn. Scale bar, 50 µm. **(B)** Bright field images of ~500 µm long 5$^{th}$ leaf primordia (left panel) and ~1700 µm long 2$^{nd}$/3$^{rd}$ leaf (right panel) expressing *GUS*-based reporter transgene for *MIR319C* or *CUC2* promoter as indicated. Black outlines of the leaf primordia/ leaves are hand drawn. **(C)** Schematic representation of the domains (colored regions) selected for 'in silico' screening for TFs specific to SAM and leaf primordia used in the Y1H screen. **(D)** Outline of the approach used to select a subset of 378 TF candidates for the Y1H screen with 2736 bp *MIR319C* upstream regulatory region (URR) as bait. **(E)** Images of yeast growth harboring *MIR319C* URR and corresponding NAC domain TF preys as indicated, grown in the presence of inhibitory concentration (25 and 30 mM) of the *HIS3* inhibitor 3-amino-1,2,4-triazole (3-AT). Negative controls including the empty vector (EV) are labeled red.

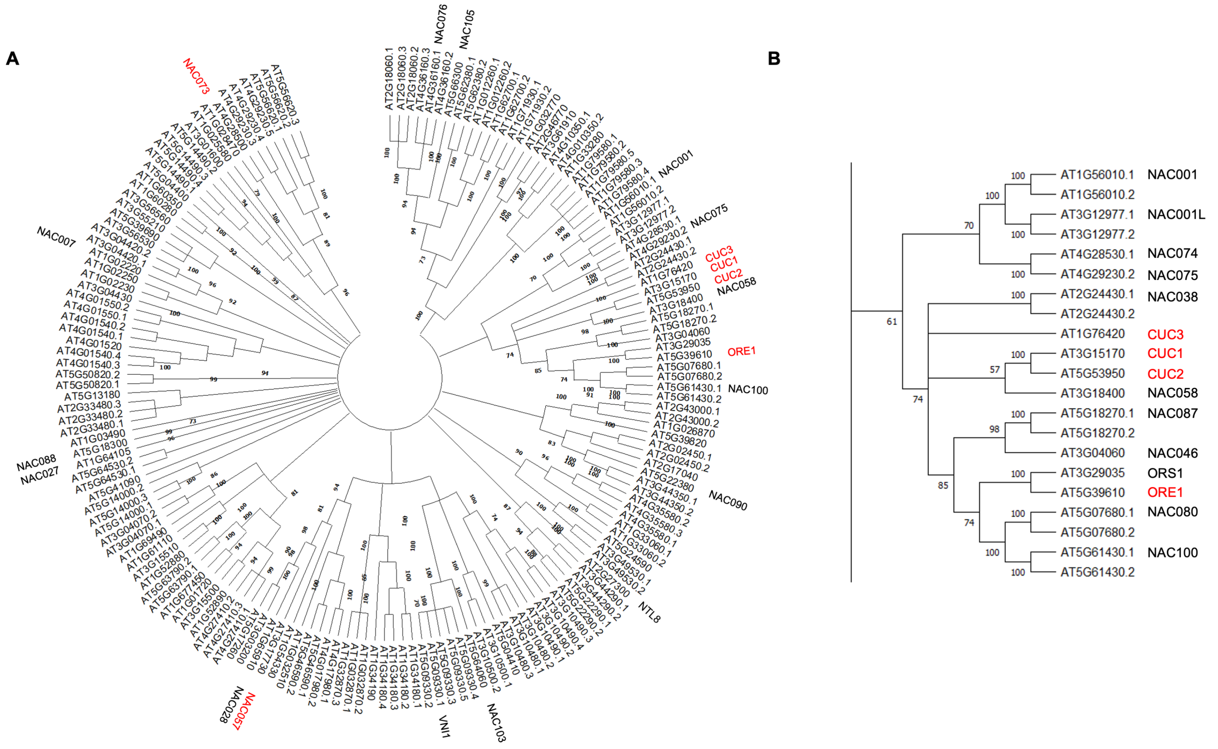


**Figure S2 (Supports Fig. 1). Phylogenetic tree of Arabidopsis NAC proteins. (A)** Maximum likelihood and JTT matrix model-based phylogenetic tree of 168 Arabidopsis NAC proteins (AtNACs). The 20 AtNACs used for the Y1H assay with truncated *MIR319C* promoter fragments (**Fig. 1C**) are labelled next to their corresponding TAIR IDs. AtNACs that bound specifically to the distal region (-2736 bp to -1866 bp) of the *MIR319C* promoter are colored in red. A part of the phylogenetic tree in **(A)** consisting of AtNAC sub families 1a (NAM/CUC3) and 1b (NAC001) is magnified in **(B)**.

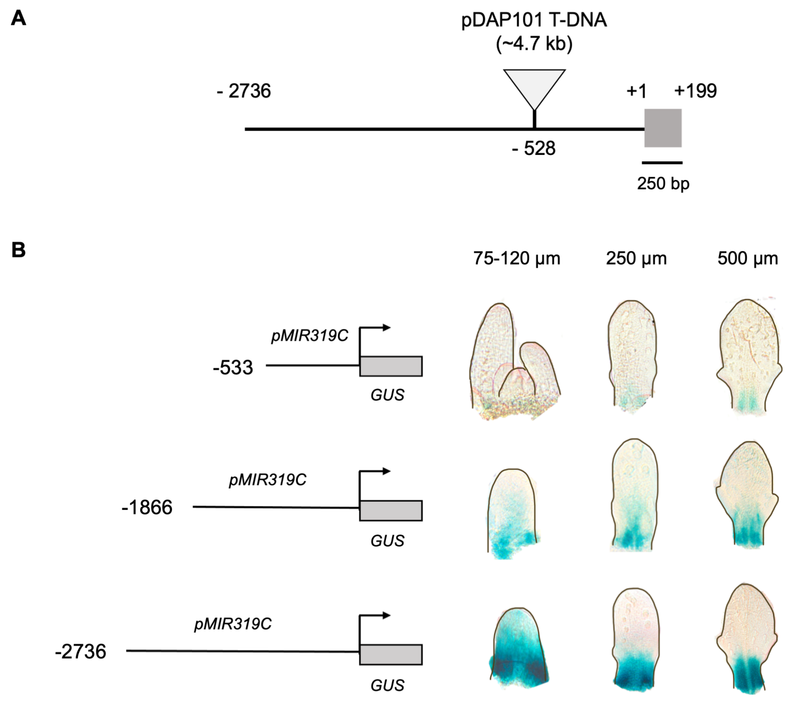


**Figure S3 (Supports Fig. 2). Establishing a new *MIR319C* mutant line. (A)** Schematic of the *miR319c-1* locus carrying 4.7 kb T-DNA insertion (indicated by inverted grey triangle) at the 528$^{th}$ nucleotide position in the *MIR319C* upstream regulatory region. The 199 bp pre-*MIR319C* region is indicated by grey box. **(B)** Bright field images of 75-120 µm long, cleared 7$^{th}$/8$^{th}$ leaf primordia (left column), 250 µm long 6$^{th}$ leaf primordia (middle column), and 500 µm long 5$^{th}$ leaf primordia (right column) expressing *GUS*-based truncated (-533 bp, -1866 bp) or full-length (-2736 bp) *MIR319C* promoter reporter transgene (*pMIR319C*) as indicated in the schematic (left side of the panels). Black outlines of the SAM and leaf primordia are hand drawn. Grey box indicates *GUS* region, and arrow indicates start of the *GUS* coding sequence (*CDS*).

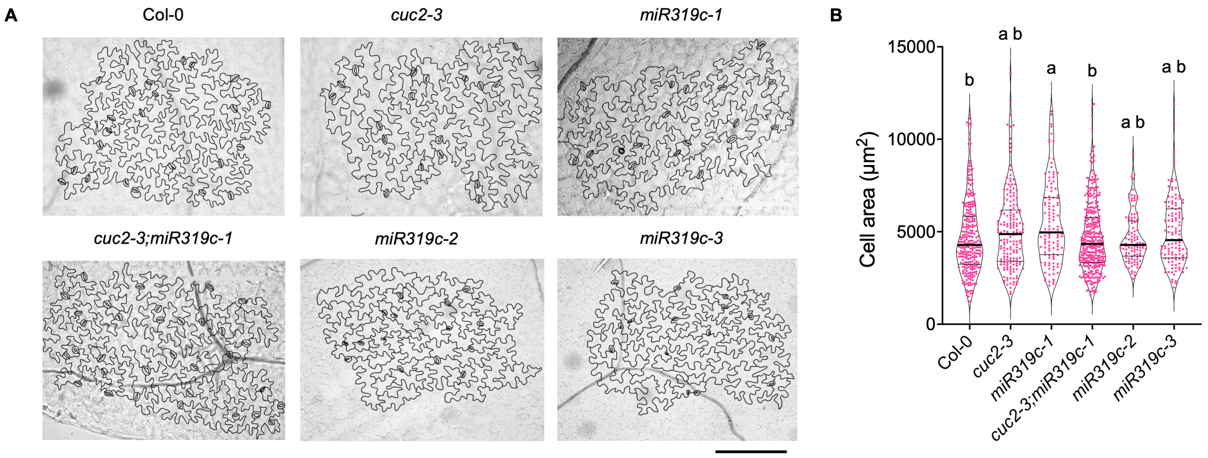


**Figure S4 (Supports Fig. 2). Pavement cell area in mutants with altered *CUC2* and *MIR319C* level.** **(A)** Representative images for pavement cells (outlines highlighted) of mature first pair of leaves from 25-day old plants of the indicated genotypes. Stomata are outlined in black. Scale bar, 200 µm. **(B)** Distribution of pavement cell area of mature first pair of leaves from 25-day old plants of the indicated genotypes. N, 105-333 cells from 3-4 leaves. Significant differences among the samples are indicated by lower-case letters. $p<0.01$; one-way ANOVA followed by Dunn's multiple comparison test was performed to determine the significant differences among samples.

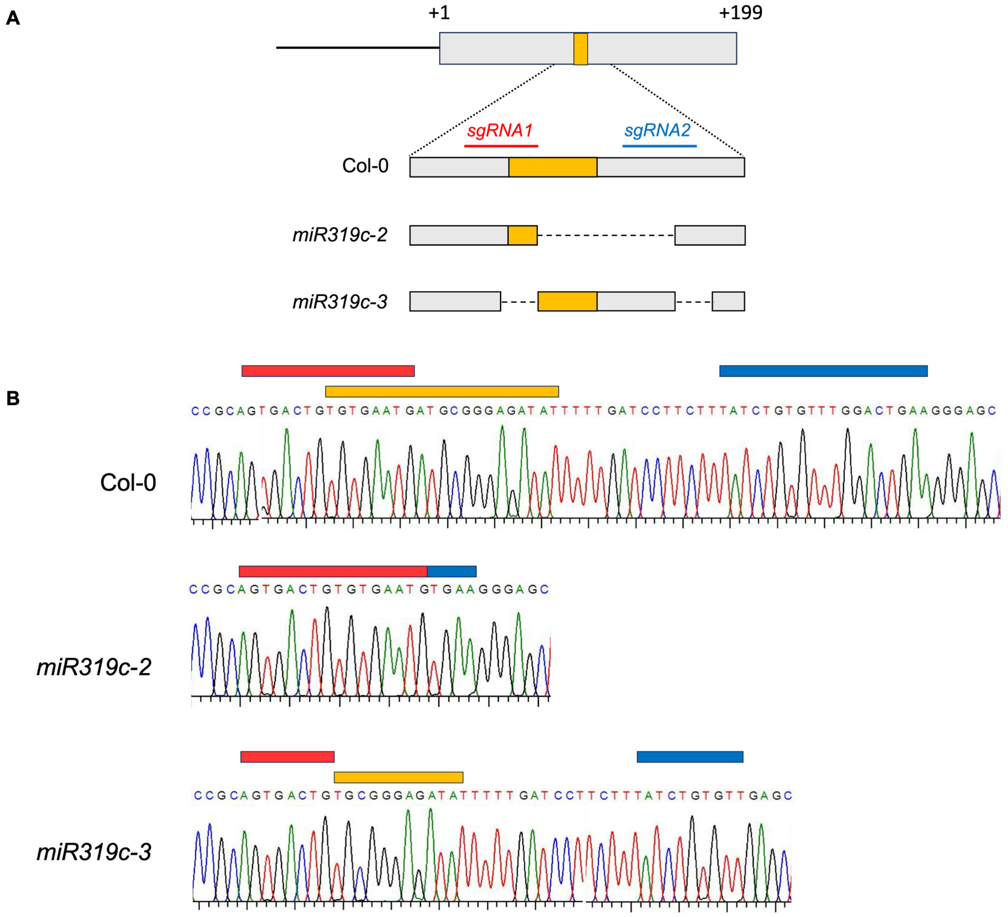


**Figure S5 (Supports Fig. 2). Generation of CRISPR-based *miR319c* null mutants. (A)** Schematic representation of *MIR319C* locus with the 199 bp pre-*MIR319C* region indicated by grey box and mature miR319c region indicated by yellow box within it. A part of the *MIR319C* locus flanked by two guide RNAs, and the mature miR319c in wild-type and *miR319c* mutants are expanded below. Dashed lines indicate deleted regions in *miR319c* mutants. **(B)** Sequence chromatogram of the *MIR319C* locus spanning guide RNA1 (horizontal red bar), mature miR319c (horizontal yellow bar) and guide RNA2 (horizontal blue bar) in Col-0 and *miR319c* mutants.

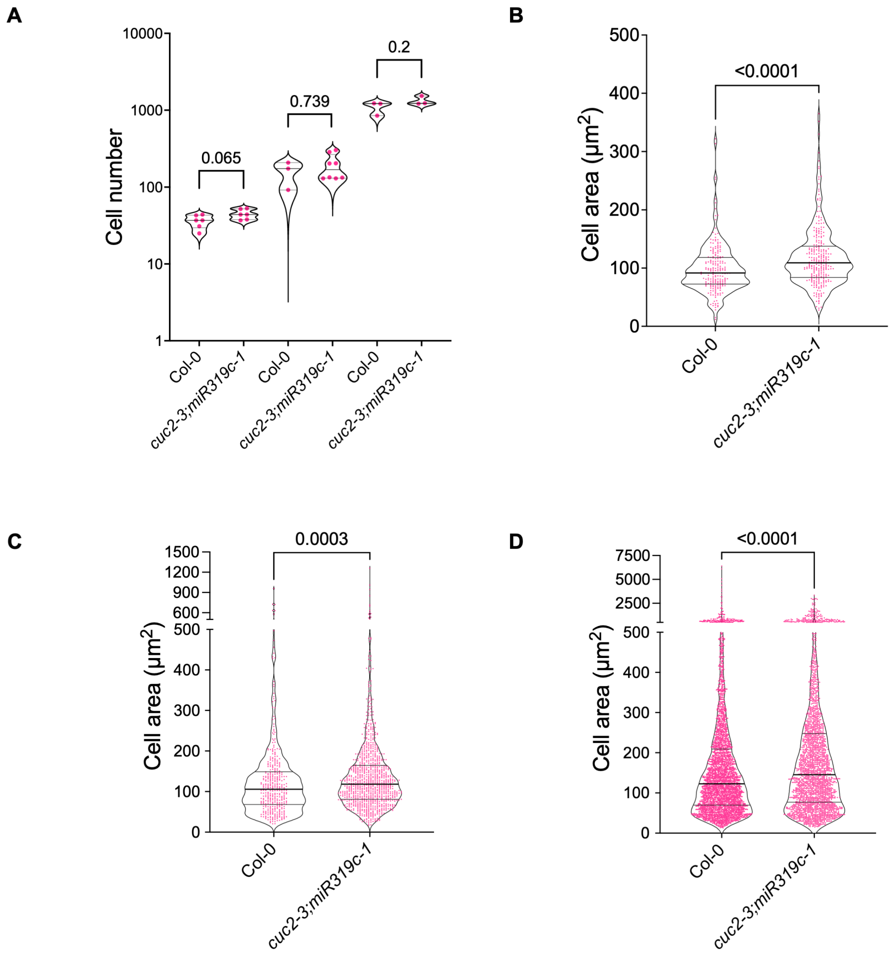


**Figure S6 (Supports Fig. 3). Number and area of pavement cells during early growth stages in *cuc2-3;miR319c-1*. (A)** Distribution of pavement cell number in leaf primordia at 1, 3, and 5 days after initiation (DAI) of Col-0 and *cuc2-3;miR319c-1*. N, 3-8 leaves. Differences among samples are indicated by *p*-values on top of the comparisons; Mann Whitney U test was used. **(B-D)** Distributions of pavement cell area of leaf primordia at 1 (**B**), 3 (**C**) and 5 (**D**) days after initiation (DAI) of Col-0 and *cuc2-3;miR319c-1*. N, 215-269 cells from 3-4 leaves (**B**), 380-955 cells from 2-3 leaves (**C**), or 2435-3280 cells from 2-3 leaves (**D**). Differences among samples are indicated by *p*-values on top of the comparisons; Mann Whitney U test was used.

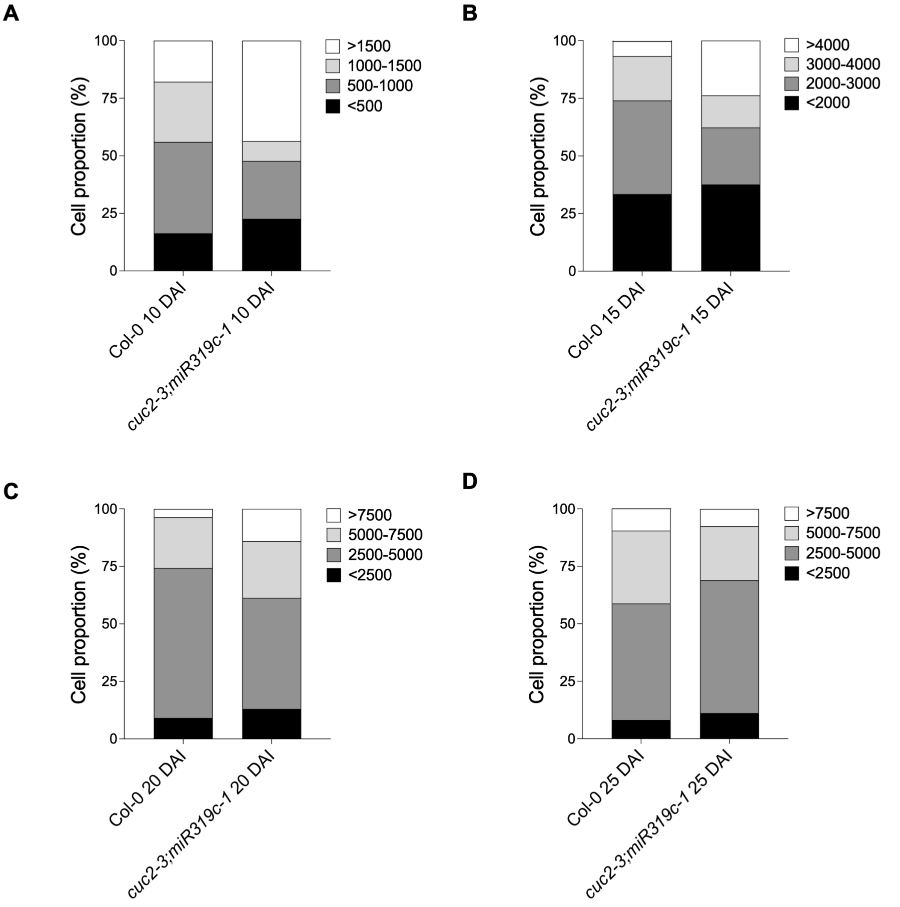


**Figure S7 (Supports Fig. 3). Cell size distribution in wildtype and *cuc2-3;miR319c-1* mature leaves. (A-D)** Stacked bar plots representing the proportion of cells (in %) with areas within the indicated size ranges (in $\mu m^2$) at 10 **(A)**, 15 (**B**), 20 **(C)** or 25 **(D)** DAI, respectively.

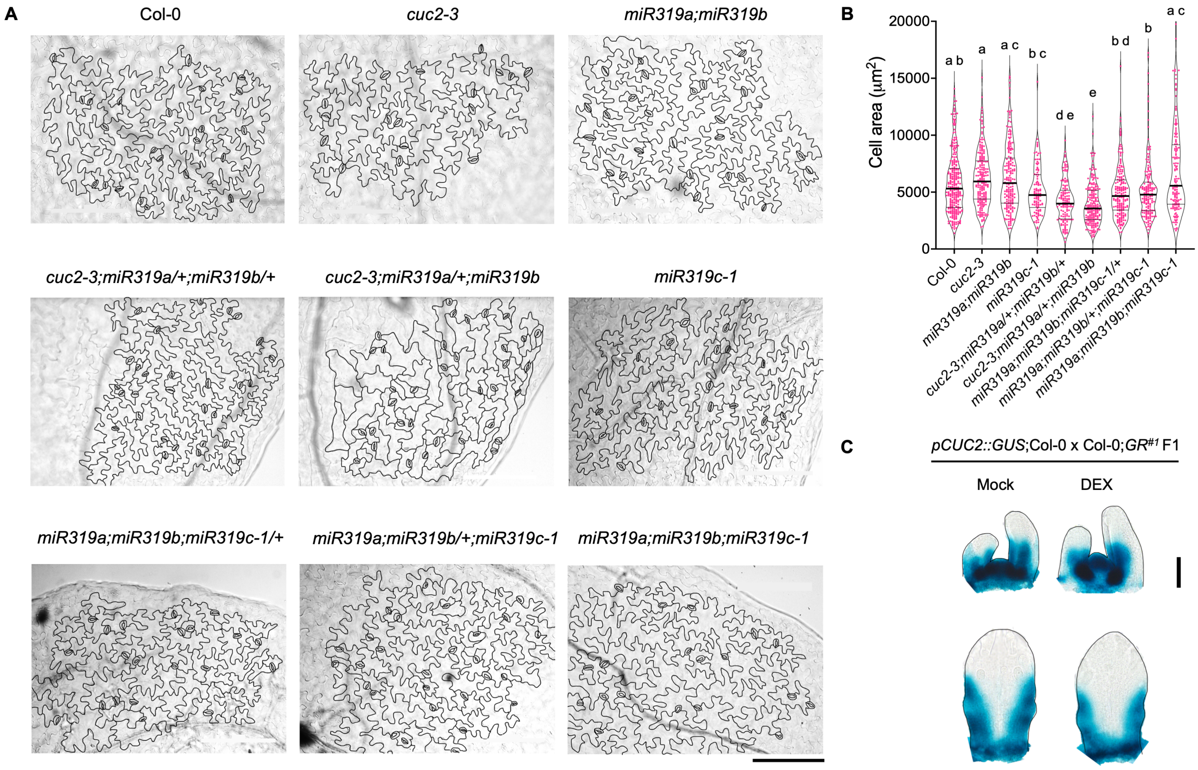


**Figure S8 (Supports Figs. 4, and 6). Pavement cell area in *cuc2;miR319* and *miR319* mutants. (A)** Representative images of pavement cells (outlines highlighted) of mature first pair of leaves from 25-day old plants of the indicated genotypes. Scale bar, 200 µm. **(B)** Distribution of pavement cell area of mature first pair of leaves from 25-day old plants of the indicated genotypes. N, 105-280 cells from 3-4 leaves. Significant differences among the samples are indicated by lower-case letters. $p<0.05$; one-way ANOVA, followed by Dunn's multiple comparison test were performed to determine the significant differences among samples. **(C)** Bright field images of shoot apices (top panel) and 6$^{th}$ leaf primordia (bottom panel) from 10-day old ethanol (Mock) or DEX-treated Col-0;*GR*$^{\#1}$ seedlings expressing *pCUC2::GUS* transgene. Black outlines of the SAM and leaf primordia are hand drawn. Scale bar, 50 µm.

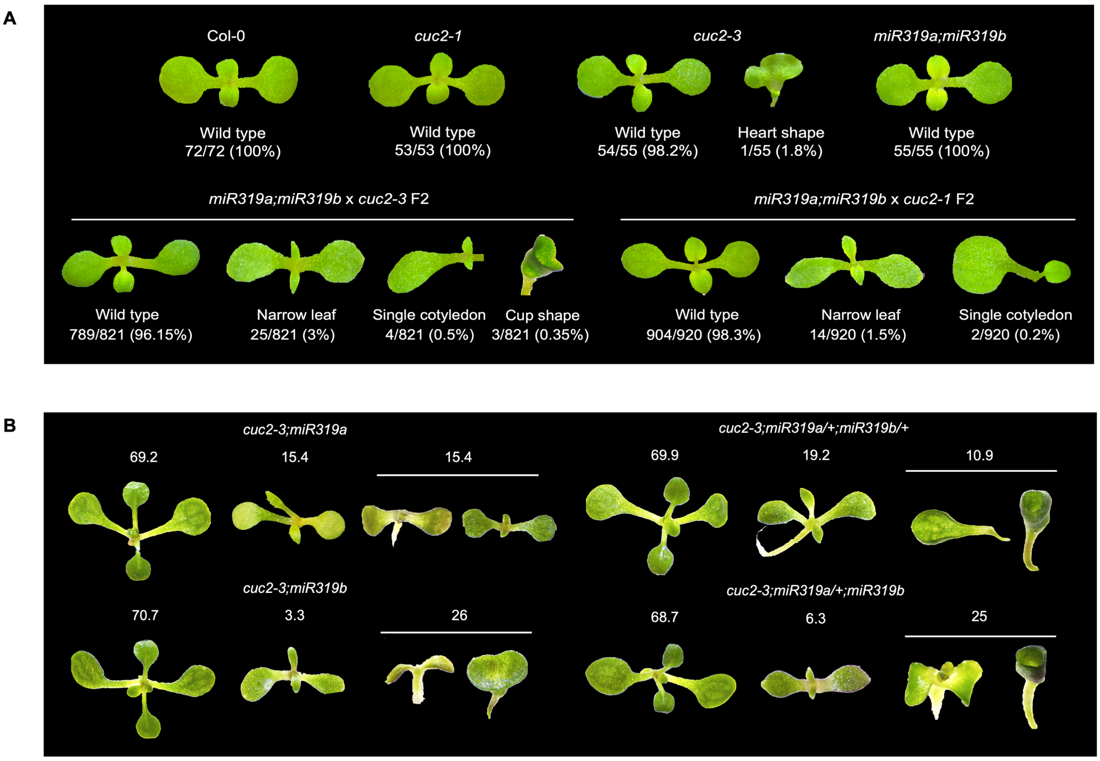


**Figure S9 (Supports Fig. 4). Leaf phenotype of mutants with altered *CUC2* and *MIR319* level. (A)** Representative images of 6-day old seedlings for each of the three categories of phenotypes, i.e., wild-type, narrow leaf, and cotyledon defects observed in the *miR319a;miR319b* x *cuc2-3* F2 and *miR319a;miR319b x cuc2-1* F2 progeny populations. Numbers below the images denote the proportion of seedling population (in %) displaying the indicated phenotype. **(B)** Representative images of 9-day old seedlings for each of the three categories of phenotypes, i.e., wild-type, narrow leaf, and seedling arrest/cotyledon fusion observed in the *cuc2;miR319a;miR319b* mutant combinations. Numbers above the images denote the proportion (in %) of seedling populations displaying the indicated phenotype.
